# Favorable histo-molecular remodeling of pancreatic ductal adenocarcinoma after Total Neoadjuvant Therapy including Stereotactic Body Radiotherapy

**DOI:** 10.1101/2024.04.30.591890

**Authors:** Christelle Bouchart, Oier Azurmendi Senar, Julie Navez, Laurine Verset, Anaïs Boisson, Matthieu Hein, Kosta Stosic, Eric Quertinmont, Vjola Tafciu, Shulin Zhao, Léo Mas, Nicky D’Haene, Dirk Van Gestel, Luigi Moretti, Ilse Rooman, Vincent Detours, Jean-Baptiste Bachet, Pieter Demetter, Karen Willard-Gallo, Rémy Nicolle, Tatjana Arsenijevic, Jean-Luc Van Laethem

## Abstract

Pancreatic ductal adenocarcinoma (PDAC) remains one of the deadliest tumors with slow progress in systemic therapies due to its peculiar and resistant tumor microenvironment. Inclusion of isotoxic high-dose stereotactic body radiation therapy (iHD-SBRT) into a total neoadjuvant strategy (TNT) is promising for the treatment of localized PDAC. However, the histo-molecular effects of iHD-SBRT are still poorly explored. In this study, we have shown that TNT, associating FOLFIRINOX [FFX] followed by iHD-SBRT, leads to significant and long-lasting remodeling of PDAC, affecting its stromal, metabolic, and molecular features. Contrary to FFX alone, TNT is able to enrich tumors with Classical and Inactive stromal signatures associated with better prognosis. Furthermore, iHD-SBRT seems capable to counteract several of the detrimental modulatory effects induced by FFX such as Epithelial-to-Mesenchymal Transition or angiogenesis. Additionally, we identified inflammatory cancer-associated fibroblasts signatures as an important prognostic factor. This work provides new rationale to sequentially combine FFX with iHD-SBRT and suggests new pathways that can be targeted in combination with a TNT.

**GRAPHICAL ABSTRACT:** 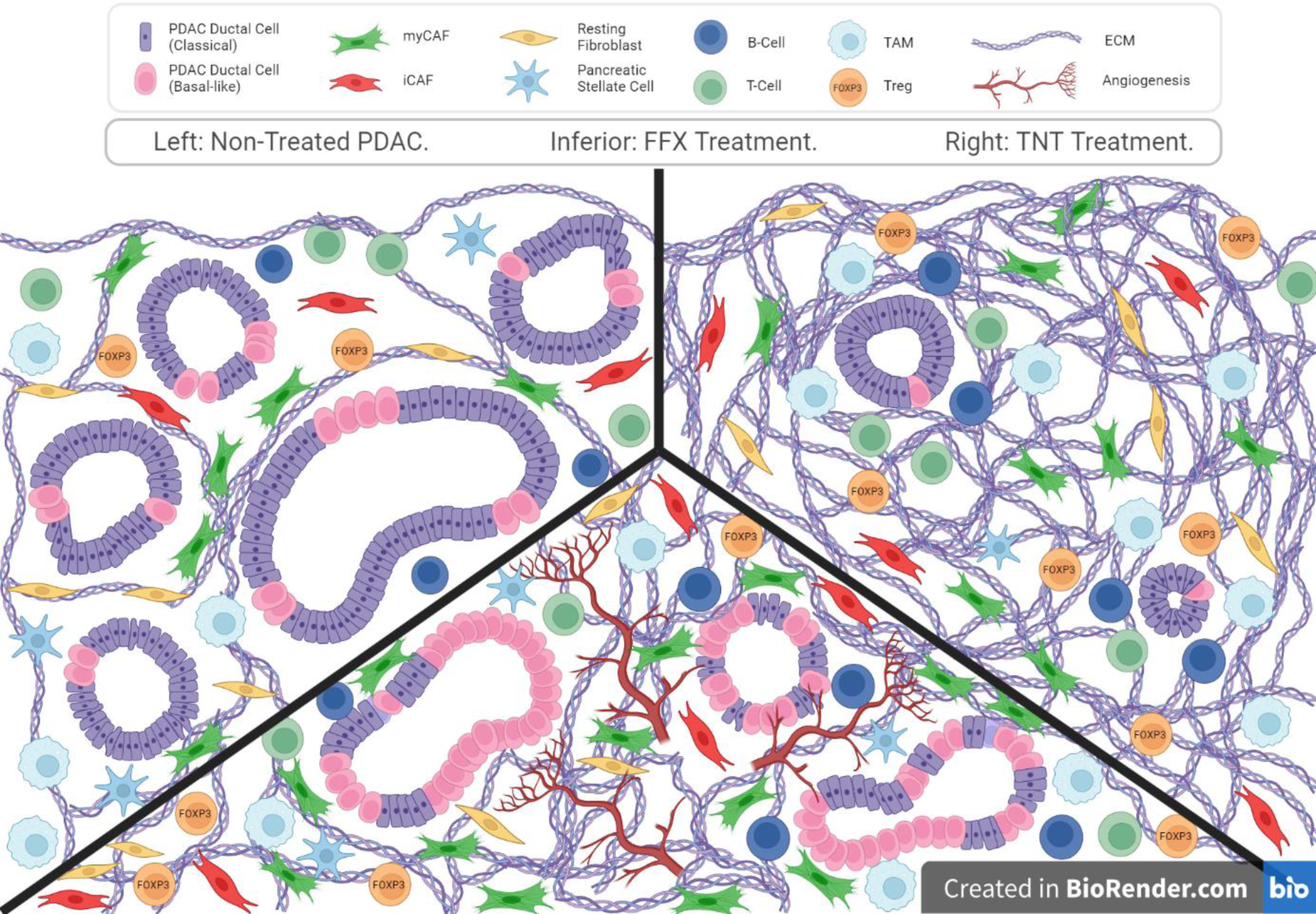

## INTRODUCTION

As of today, pancreatic ductal adenocarcinoma (PDAC) remains one the deadliest tumors, with a 5-year survival rate of less than 12%. [1] Despite recent improvements in the therapeutic arsenal with the introduction of more active multi-agent chemotherapy (FOLFIRINOX [FFX], Gemcitabine / nab-paclitaxel or NALIRIFOX), progress in systemic therapies for PDAC has been slow compared to other cancers. [2–4] While many clinical trials have explored the efficacy of immune checkpoint inhibitors (ICIs), cancer vaccines, or targeted therapy, these have not led to major changes in clinical practice. [5,6] The difficulty in obtaining concrete oncological benefits in these clinical trials stems largely from the peculiar tumor microenvironment (TME) of PDAC, which provides many paths of resistance and aggressiveness. [5,6] Therefore, it is crucial to better comprehend the complexity and the crosstalk mechanisms involved, as well as to improve our understanding of how modern therapies currently used in clinical practice influence the modulation of the TME.

Neoadjuvant therapy is a rapidly growing strategy for non-metastatic PDAC patients, although the exact sequence to use remains to be determined. [7] The FFX regimen is currently the preferred chemotherapy used in the neoadjuvant setting by many centers due to the results of several trials in metastatic and non-metastatic patients showing a significant superiority in survival compared to gemcitabine alone, as well as a safe and active profile in neoadjuvant phase II trials. [2, 8–11] The addition of (nearly) ablative stereotactic body radiation therapy (SBRT) to multi-agent chemotherapy in the neoadjuvant setting as a total neoadjuvant therapy (TNT) may offer several advantages over conventional chemoradiotherapy (CRT). These include notably the capacity to deliver more easily and rapidly a higher biologically effective dose (BED) to the tumor, associated with improved survival outcomes, as well as a shorter break of full-dose chemotherapy. [7, 12–14] Several studies reported promising results and an increasing number of (randomized) phase II clinical trials are currently exploring this question, including ours (STEREOPAC trial – NCT05083247). [7,14–18] However, if radiation therapy is able modulate the TME, the impact of modern high-dose SBRT (> 35Gy in 5 fractions) on the immune components and other molecular features is still poorly known in PDAC. A better understanding of these modulations may pave the way for the development of molecularly oriented combination trials with immune and/or targeted therapies as well as stratified treatment strategies, which are urgently needed in PDAC. The identification of molecular subtypes in PDAC has gained a lot of interest in the last decade and it is now clearer that these molecular signatures have the potential to lead to better selection of patients, the prediction of the response to treatments and therefore, the development of individualized treatments. [19–26] While the relationship between molecular subtypes and chemotherapy is progressively explored, little is known regarding RT and to our knowledge, nothing for high-dose SBRT nor its inclusion into a TNT sequence.

In this study, we aimed to characterize for the first time in PDAC the molecular subtypes, transcriptomic profiles and immuno-modulations following FFX alone or in a TNT including isotoxic high-dose SBRT (iHD-SBRT). We hypothesized that iHD-SBRT can sustainably modify the molecular and transcriptional profiles in PDAC, shedding light on key cells and pathways involved and leading to a better understanding of the respective contribution and complementarity of a TNT.

## RESULTS

### Patients characteristics and outcomes

A total cohort of 124 retrospectively collected patients treated for localized PDAC and surgically resected between 2011 and 2020 was assessed for eligibility. Sixty-five patients were initially included, but fifteen were subsequently excluded, as they did not meet the RNA sequencing (RNAseq) quality check criteria. Finally, RNAseq data from 50 PDAC patients were considered for this study. This cohort comprises: 1/ Seventeen patients in the non-neoadjuvant (No_NAT) group; 2/ Seventeen patients in the FFX group and 3/ Sixteen patients in the TNT group (FFX followed by iHD-SBRT before surgery). The methodology workflow of the study is described in the CONSORT-like clinic-molecular diagram in **Fig. 1**. In the TNT cohort, the patients underwent an oncological surgical resection at a median time of 44 days (31 - 70 days) after iHD-SBRT, and this group included significantly more locally advanced patients. No significant difference in median overall survival (OS) or median disease-free survival (DFS) were observed between the three groups. However, we noted that the 1-year DFS was significantly improved in the TNT cohort (TNT vs FFX vs No_NAT: 87.5 vs 70.6 vs 41.2%, respectively, p=0.017) (**Supplementary Fig. 1**). The main clinico-pathological characteristics of the included patients are summarized in **Table 1**.

**Figure 1.**
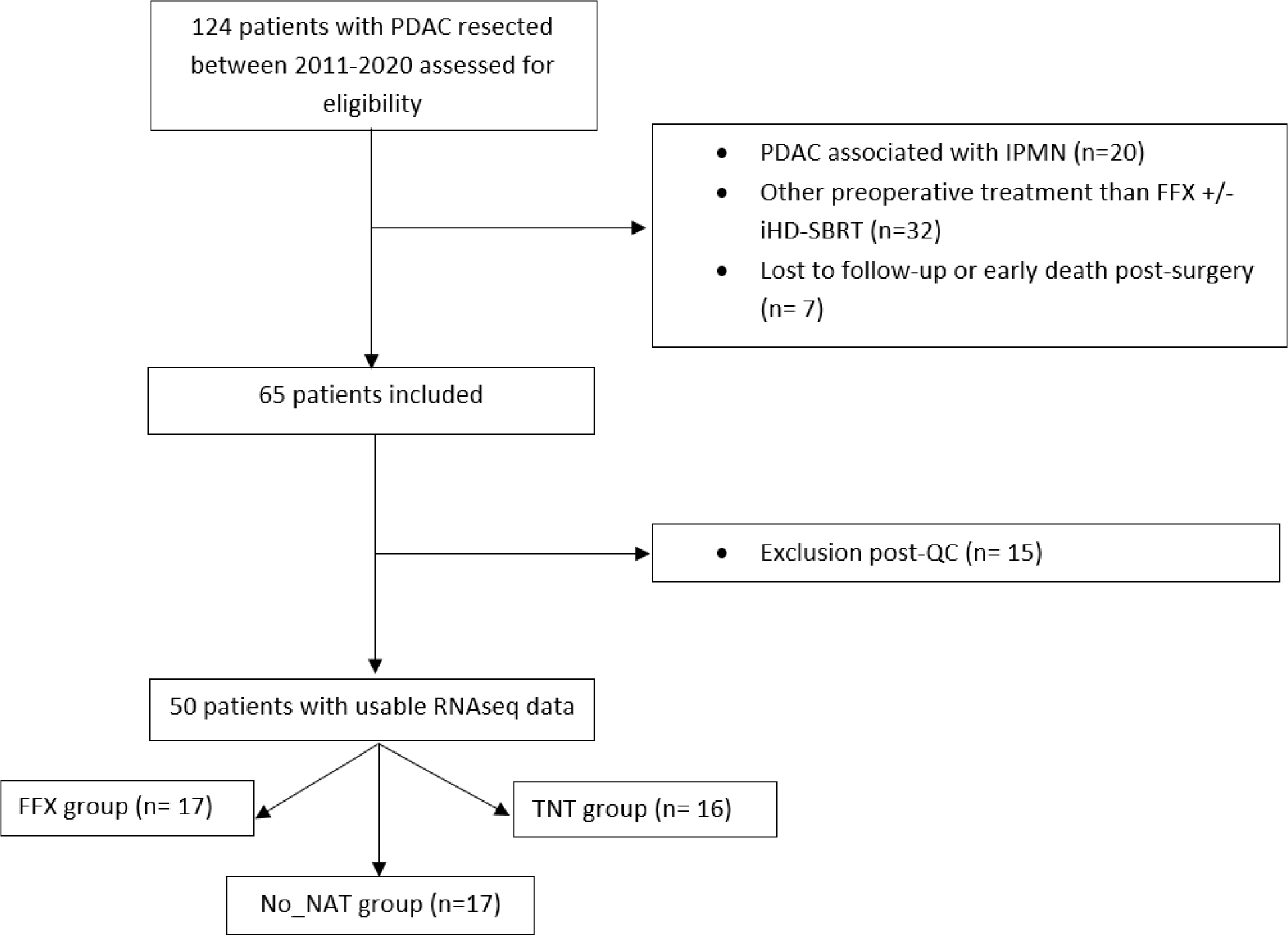
CONSORT-like workflow description of the PDAC cohort. Detailed description of the selection process of the patients and samples cohort. PDAC: pancreatic ductal adenocarcinoma; IPMN: intraductal papillary mucinous neoplasm; FFX: FOLFIRINOX; iHD-SBRT: isotoxic high-dose stereotactic body radiotherapy; QC: quality control; TNT: total neoadjuvant treatment (FFX + iHD-SBRT); No_NAT: no neoadjuvant treatment group.

**Table 1.** Main characteristics and outcomes of the studied cohort. No_NAT= non-treated; FFX= FOLFIRINOX; TNT= total neoadjuvant treatment (FFX + iHD-SBRT); FU = follow-up; DFS= disease free survival; OS= overall survival

|  | Whole cohort<br>(n=50) | No_NAT<br>(n=17) | FFX<br>(n=17) | TNT<br>(n=16) | P-value |
| --- | --- | --- | --- | --- | --- |
| **Age (years)** | 66.8 (57.6 – 69.8) | 69.1 (60.5 – 70.9) | 64.7 (57.6 – 66.8) | 67.0 (53.2 – 70.4) | 0.126 |
| **Gender (%)** |  |  |  |  | 0.644 |
| Male | 56.0 | 47.1 | 58.8 | 62.5 |  |
| Female | 44.0 | 52.9 | 41.2 | 37.5 |  |
| **Clinical staging TNM 8<sup>th</sup> ed. (%)** |  |  |  |  | <0.001<sup>a,b</sup> |
| IA | 10.0 | 29.4 | 0.0 | 0.0 |  |
| IB | 32.0 | 58.8 | 29.4 | 6.3 |  |
| IIA | 0.0 | 0.0 | 0.0 | 0.0 |  |
| IIB | 30.0 | 11.8 | 35.3 | 43.7 |  |
| III | 28.0 | 0.0 | 35.3 | 50.0 |  |
| **CA19.9 values at diagnosis (kU/L)** | 49.3 | 28.5 | 58.0 | 101.9 | 0.265 |
| **Tumor diameter (mm)** | 28.0 (22.0 – 35.0) | 22.0 (17.0 – 25.0) | 30.0 (26.0 – 35.0) | 37.5 (27.6 – 46.0) | <0.001<sup>a,b</sup> |
| **Primary site (%)** |  |  |  |  | 0.571 |
| Head/uncus/isthmus | 88.0 | 94.1 | 82.4 | 87.5 |  |
| Body/tail | 12.0 | 5.9 | 17.6 | 12.5 |  |
| **Resection status (%)** |  |  |  |  | <0.001<sup>a,b</sup> |
| Resectable | 46.0 | 94.1 | 35.3 | 6.2 |  |
| Borderline resectable | 40.0 | 5.9 | 52.9 | 62.5 |  |
| Locally advanced | 14.0 | 0.0 | 11.8 | 31.3 |  |
| **Number of neoadjuvant FFX cycles received** | 6 (5 – 8) | / | 6 (4 – 8) | 7 (6 – 8) | 0.378 |
| **Pathological staging TNM 8<sup>th</sup> ed. (%)** |  |  |  |  | 0.697 |
| IA | 10.0 | 5.9 | 17.7 | 6.2 |  |
| IB | 18.0 | 17.6 | 17.6 | 18.8 |  |
| IIA | 4.0 | 0.0 | 0.0 | 12.5 |  |
| IIB | 26.0 | 35.3 | 17.6 | 25.0 |  |
| III | 34.0 | 35.3 | 35.3 | 31.2 |  |
| IV | 8.0 | 5.9 | 11.8 | 6.3 |  |
| **Differentiation grade (%)** |  |  |  |  | 0.006<sup>b</sup> |
| Good | 12.2 | 0.0 | 23.5 | 13.3 |  |
| Intermediate | 42.9 | 29.4 | 29.4 | 73.3 |  |
| Poor | 44.9 | 70.6 | 47.1 | 13.4 |  |
| **Adjuvant chemotherapy received (%)** |  |  |  |  | 0.492 |
| No | 16.0 | 11.8 | 11.8 | 25.0 |  |
| Yes | 84.0 | 88.2 | 88.2 | 75.0 |  |
| **FU, median [IC95%] (months)** | 29.5 (19.0 – 50.0) | 24.1 (18.4 – 52.6) | 36.9 (19.0 – 47.6) | 28.3 (21.6 – 49.9) | 0.821 |
| **DFS, median [IC95%] (months)** | 17.5 (12.8 – 21.6) | 10.0 (5.0 – 20.4) | 17.7 (7.0 – 35.6) | 20.6 (15.6 – 27.3) | 0.496 |
| **1-year DFS (%)** | 66.0 | 41.2 | 70.6 | 87.5 | 0.017<sup>b</sup> |
| **OS, median [IC95%] (months)** | 31.8 (24.1 – 47.6) | 24.1 (17.3 – 54.4) | 38.5 (18.5 – 97.5) | 32.3 (22.4 – 75.5) | 0.558 |

### iHD-SBRT following chemotherapy induction with FFX is able to reverse several of the main unfavorable transcriptome alterations induced by FFX in PDAC

To examine the biological functions of the identified differentially expressed genes (DEGs) between different groups, we performed the gene ontology (GO) functional annotation describing genes and their associations according to three ontology categories (molecular function, cellular component and biological process) (**Fig. 2** and **Supplementary Fig. 2**). [27] In the GO analyses, the FFX group, compared to the No_NAT samples, demonstrated a significant positive enrichment in mitotic cell cycle arrest, extracellular matrix (ECM), transcriptional activity (including histone demethylation), regulation of glucose transport, as well as for regulation of angiogenesis, the vascular endothelial growth factor (VEGF) signaling pathway and epithelial to mesenchymal transition (EMT). Conversely, when iHD-SBRT was added to FFX in the therapeutic strategy (TNT vs FFX group), we interestingly observed a significant negative enrichment in glucose transport, angiogenesis-related items, as well as ECM assembly and EMT process. Furthermore, the TNT group showed significant positive enrichment scores notably related to mitochondrial activity, glutathione biosynthetic process and apoptotic cell clearance, while a reduced level of items was detected related to cell adhesion, cell migration (including for fibroblasts), ECM organization and cellular response to TGFβ stimulus.

**Figure 2.**
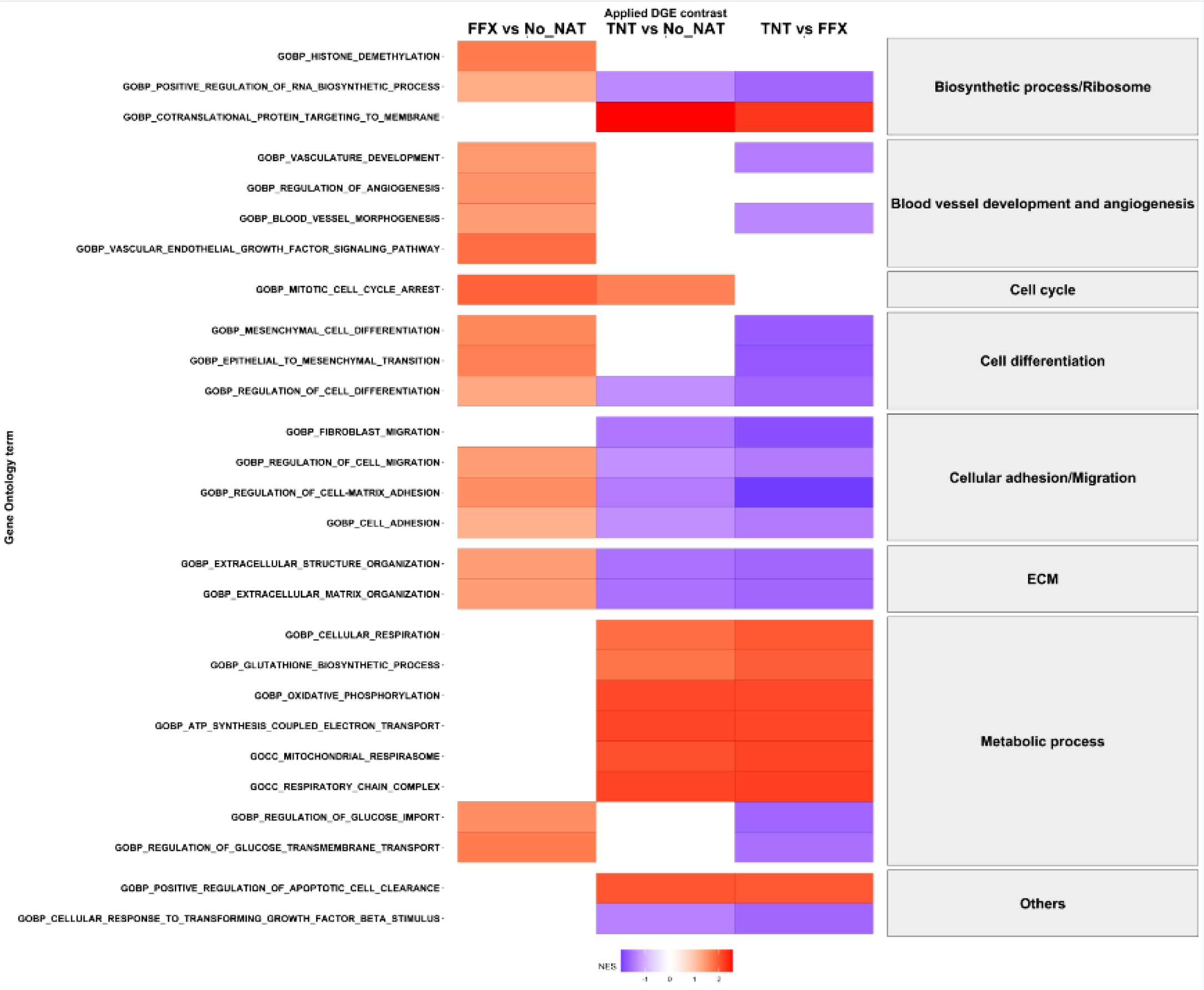
Differential enrichment via gene set enrichment analysis of the Gene Ontology (GO) terms following neoadjuvant treatments. Selected Gene Set Enrichement Analysis (GSEA) results of the Gene Ontology (GO) terms, grouped according to the biological function, with differential gene expression comparisons between the three groups. FFX: FOLFIRINOX; No_NAT: no neoadjuvant treatment group; TNT: total neoadjuvant treatment (FFX + iHD-SBRT); ECM: extracellular matrix.

GO and Molecular signatures database (MSigDB) canonical pathways and, consistently, mitochondrial activity, glutathione metabolism and ribosomal pathways were significantly enriched post-iHD-SBRT (TNT vs No_NAT only). Additionally, when single-nucleus signatures from Hwang *et al*. were applied, significant enrichments were found for the ribosomal biogenesis whereas TNF/NF-kB signaling exhibited reduced level after iHD-SBRT (**Fig. 3**). [25]

**Figure 3.**
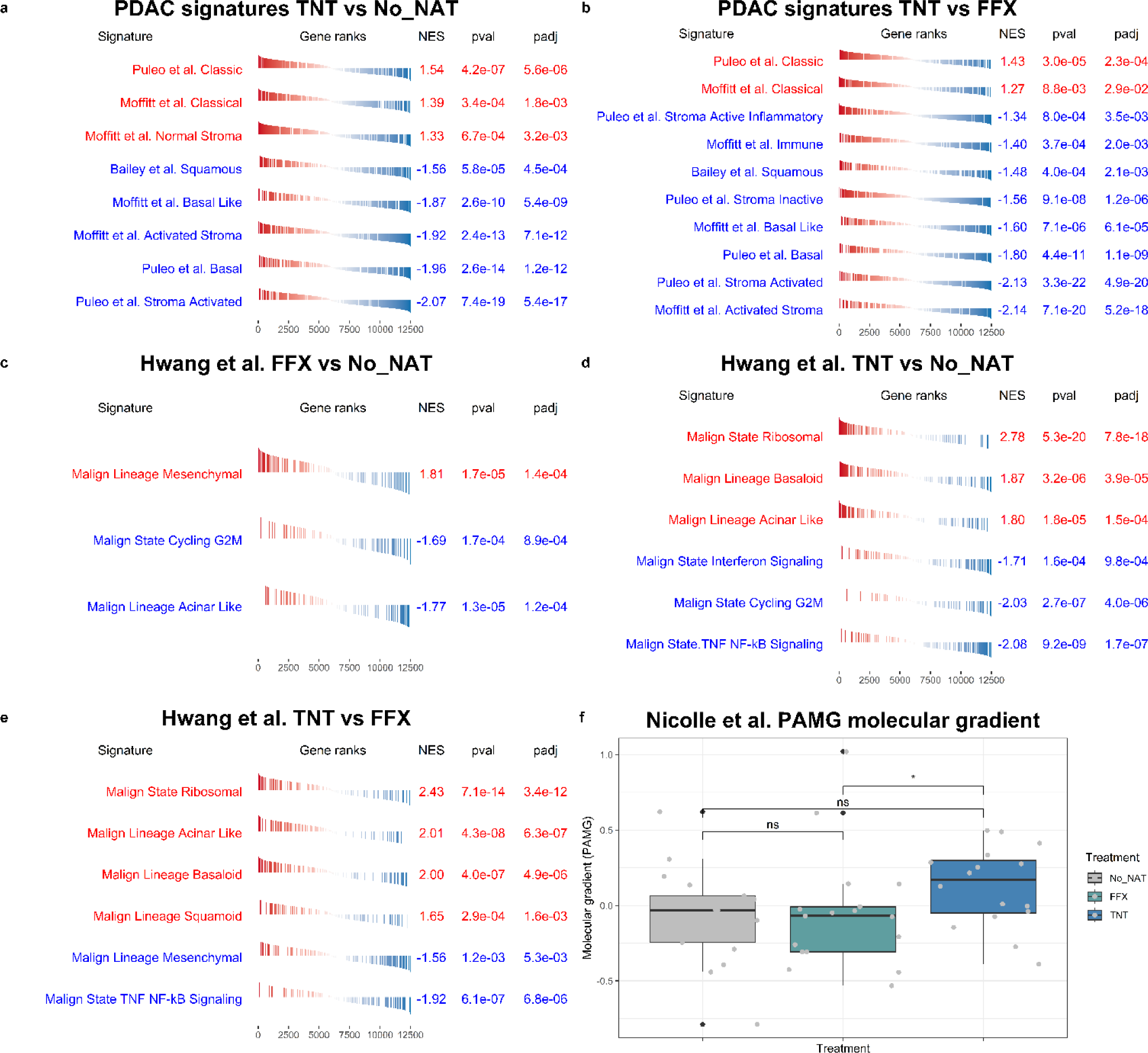
Enrichment analyses of the tumoral molecular subtypes and cell types between the three groups. **(a,b)** Normalized Enrichment Score (NES) after PDAC subtype RNA signatures enrichment analysis showing significantly higher NES for the “Classical” subtypes and decreased “Basal” subtypes in TNT group vs No_NAT **(a)** and TNT vs FFX **(b)** through the main transcriptomic PDAC classifications. **(c,d,e)** NES after GSEA of Hwang *et al.* signatures obtained with single nucleus RNA-seq. Differential gene expression comparison between FFX vs No_NAT group **(c)**, TNT vs No_NAT group **(d)** and TNT vs FFX group **(e)**, showing a significant enrichment in the Mesenchymal subtype in FFX samples whereas an enrichment of Basaloid subtype, associated with favorable prognosis, is observed in the TNT group. **(f)** A continuous gradient of PDAC pre-existing classifications, the pancreatic adenocarcinoma molecular gradient (PAMG), was applied on the whole cohort, showing a significantly higher molecular gradient PAMG score (p=0.049) in favour of the TNT group compared to FFX group. FFX: FOLFIRINOX; No_NAT: no neoadjuvant treatment group; TNT: total neoadjuvant treatment (FFX + iHD-SBRT)

### Addition of iHD-SBRT to FFX is associated with transcriptomic signatures and PAMG score linked to better prognosis

The molecular subtype signatures from the main studies available in the field (Puleo *et al*; Moffitt *et al.*; Bailey *et al*.; Hwang *et al.* [20–22, 25]) were explored in this cohort to determine the influence of modern neoadjuvant treatments, including high-dose SBRT (**Fig. 3**). When compared to both No_NAT and FFX groups, the TNT group showed a significant enrichment in the more favorable “Classical subtype” signatures (**Fig. 3a-b**, in red). Furthermore, the addition of iHD-SBRT was also associated with a reduced level of “activated stroma” and “Basal-like subtype” signatures from all major molecular classifications, which are associated with poorer prognosis (**Fig. 3a-b**, in blue).

To get a deeper insight into the evolution of molecular subtypes’ following the two different neoadjuvant treatments, we applied the recently published single-nucleus signatures from Hwang *et al*. to our cohort. [25] The FFX group compared to No_NAT was enriched with the “Mesenchymal” signature, representing a subtype of “Basal-like” cells, and several stromal signatures associated with highly active stroma, all of which being associated with worse clinical outcomes (**Fig. 3c and Supplementary Fig. 3**). The neural-like progenitor and neudroendocrine –like programs identified in Hwang *et al*. as significantly higher post chemo-radiotherapy were not significantly enriched in our cohort (**Supplementary Data 1-2**). [25] Interestingly, when compared to both the No_NAT and FFX cohorts, the TNT group was notably significantly associated with a “Basaloid” signature, representing a particular subtype of “Basal-like” cells associated with better clinical outcomes (**Fig. 3d-e**). [25]

Finally, a continuous gradient of PDAC pre-existing classifications, the pancreatic adenocarcinoma molecular gradient (PAMG), was applied and revealed a significant favorable shift in samples treated with TNT towards a higher PAMG score compared to FFX alone. These data confirm that the TNT group is significantly enriched with the “Classical” subtype gene signatures, associated with better cell differentiation, as well as improved clinical outcomes (**Fig. 3f**). [28]

### TNT modulates the metabolic state of PDAC towards an enrichment of the cholesterogenic metabolic profile

Given that FFX alone and TNT appear to induce opposite enrichment scores regarding several transcriptional items related to metabolism, such as mitochondrial activity (including oxidative phosphorylation) and glucose import (**Fig. 2b** and **Supplementary Fig. 3**), a deeper characterization of the metabolic state was performed in our cohort using the metabolic gene signatures identified by Karasinska *et al*.. [29] In all three of our groups, the glycolytic genes were significantly associated with “Basal-like” genes, while cholesterogenic genes were with “Classical” subtype genes (Supplementary Fig. 4). Compared to FFX, TNT was associated with a significant positive enrichment score related to cholesterol biosynthesis, which correlates with favorable clinical outcomes (**Supplementary Fig. 4).** [29]

### TNT generates different modulations on the cancer associated fibroblasts (CAFs) transcriptomic signatures than FFX alone

As it was observed that both FFX and TNT had a significant transcriptomic impact on stromal signatures and ECM organization, xCell analyses were performed, revealing a significantly higher stroma score (**Fig. 4a**) and CAFs population (**Fig. 4b**) after neoadjuvant FFX compared to No_NAT. Several bulk and single-cell based CAF classifications were then tested with GSEA to observe the specific CAFs modifications induced by FFX and TNT (**Fig. 4c-h**).

**Figure 4.**
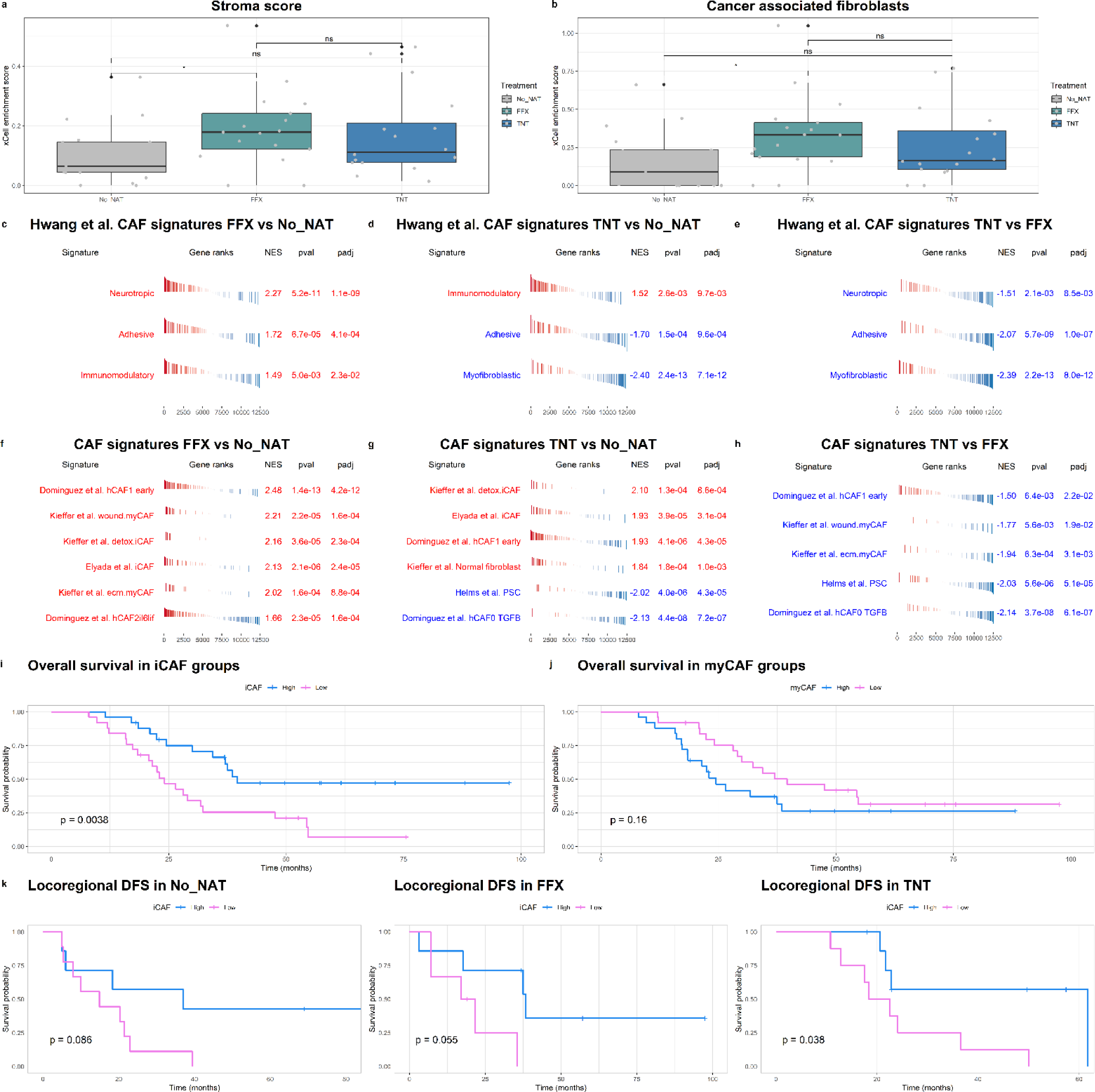
Cell type enrichment analysis of stromal and cancer associated fibroblasts (CAFs) transcriptomic signatures following neoadjuvant treatments. **(a,b)** Cell type enrichment analysis using xCell showing a significantly higher stroma score (p=0.034) **(a)** and CAFs population (p=0.039) **(b)** in FFX vs No_NAT group. **(c,d,e)** Normalized Enrichment Score (NES) after GSEA of Hwang *et al.* gene sets obtained with single nucleus RNA-seq: differential expression comparison between FFX vs No_NAT group **(c)**, TNT vs No_NAT group **(d)** and TNT vs FFX group **(e).** **(f,g,h)** NES after GSEA of state of the art CAFs gene sets: differential expression comparison between FFX vs No_NAT group **(f)**, TNT vs No_NAT group **(g)** and TNT vs FFX group **(h)**. **(i,j)** Gene set variation analysis (GSVA) was applied as a single sample classifier of different CAF subtypes defined in Elyada *et al.* to classify all the samples according to their enrichment in high and low iCAF and myCAF groups. Kaplan–Meier survival analyses were performed on high and low CAF populations. High-iCAF samples showed a significantly better overall survival (OS) compared to Low-iCAF (p=0.0038) **(i)** while no statistical difference was found for myCAFs **(j)**. (**k**) Locoregional disease free survival (LR-DFS) in the three groups stratified per high and low-iCAF samples. A significantly better LR-DFS was observed in high-iCAF in the TNT cohort (p=0.038) while a non-significant tendency has been observed for the No_NAT and FFX groups. No_NAT: untreated; FFX: FOLFIRINOX; TNT: Total neoadjuvant treatment (FFX + iHD-SBRT)

An enrichment in “Immunomodulatory” CAFs (from Hwang *et al.* [25]) and inflammatory CAFs (iCAFs) signatures was observed in both neoadjuvant cohorts compared to No_NAT. Interestingly, after FFX alone, compared to the two other groups, a significant enrichment in myofibroblastic CAFs (myCAFs) signatures, associated with worse prognosis, was observed (**Fig. 4c and f**). Furthermore, our results indicate that patients treated with TNT display fewer pancreatic stellate cells (PSCs) as well as myCAFs compared to both No_NAT and FFX groups and are enriched in “Normal Fibroblasts” signatures compared to the No_NAT samples (**Fig. 4d-e, g and h**).

### iCAFs but not myCAFs are significantly associated with better clinical outcome in No_NAT and TNT cohorts

iCAF and myCAF transcriptomics signatures from Elyada *et al.* were tested independently using the single sample classifier Gene Set Variation Analysis (GSVA) to classify all the samples according to their enrichment in high and low groups for each subtype. [30] Interestingly, in our whole cohort, iCAF-high samples had a significantly better OS than the iCAF-low group (p=0.0038) (**Fig. 4i**). In addition, the iCAF-high samples displayed a significantly better LR-DFS compared to the iCAF-low in the TNT cohort (p=0.038) (**Fig. 4k**). This observation was validated in a No_NAT external cohort (Puleo *et al.* [22]; n=309), confirming a significant difference in relapse free survival according to iCAF enrichment (p=0.041) (**Supplementary Fig. 5**). No significant differences were observed between high and low myCAF groups for DFS and OS in our cohort, nor in the Puleo *et al.* cohort (**Fig. 4j** and **Supplementary Fig. 5**). These results suggest an important potential of iCAF as a prognostic / predictive factor.

### Neoadjuvant treatments increase desmoplasia without significantly affecting tumor-infiltrating lymphocytes (TILs) except for the T helper population

To further assess the stromal characteristics of PDAC, the percentage of the tumoral area occupied by collagen was quantified through immunohistochemistry (IHC) analysis across the entire cohort. Consistent with our previously described findings, a significant increase in Collagen1A1 (COL1A1) deposition – a marker indicative of pan-fibroblast population - was observed in both neoadjuvant groups compared to the No_NAT group (68.4 vs 78.6 vs 83.27% for No_NAT vs FFX vs TNT, respectively, p<0.001) (**Fig. 5a**). Additionally, a non-significant trend towards a lower expression of αSMA (a marker associated with myCAFs) was noticed in the TNT group compared to FFX (**Fig. 5b**). Notably, despite the increase in collagen deposition in tumors treated with neoadjuvant treatments, no significant changes were observed in the expression levels of CD3 TILs as well as cytotoxic CD8+ cells, including after TNT (**Fig. 5c-d**). Regarding T-cells, only the CD4+ T helper population was significantly deceased after TNT compared to FFX and No_NAT groups (**Fig. 5e**). The B-cell CD20+ population was decreased after NAT with a significant difference observed between TNT and No_NAT groups (**Supplementary Fig. 6**). Following review by specialized GI pathologists (LV and PDM), signs of tumoral cells injury such as cell swelling and pyknotic nucleus were often observed after NAT (**Fig. 6)**. The immune infiltration including tumor infiltrating lymphocytes (TILs) did not appear to be sequestrated in the collagenous stroma after TNT, and remained present in close proximity to the remaining tumoral glands, with TILs infiltrating directly within the tumoral glands, as illustrated in **Fig. 6f**. The presence of scarce tertiary lymphoid structures (TLS) within the tumoral area was identified on consecutive H&E and CD3/CD20 dual stained slides and no significant difference was observed between the three groups (**Supplementary Fig. 6**).

**Figure 5.**
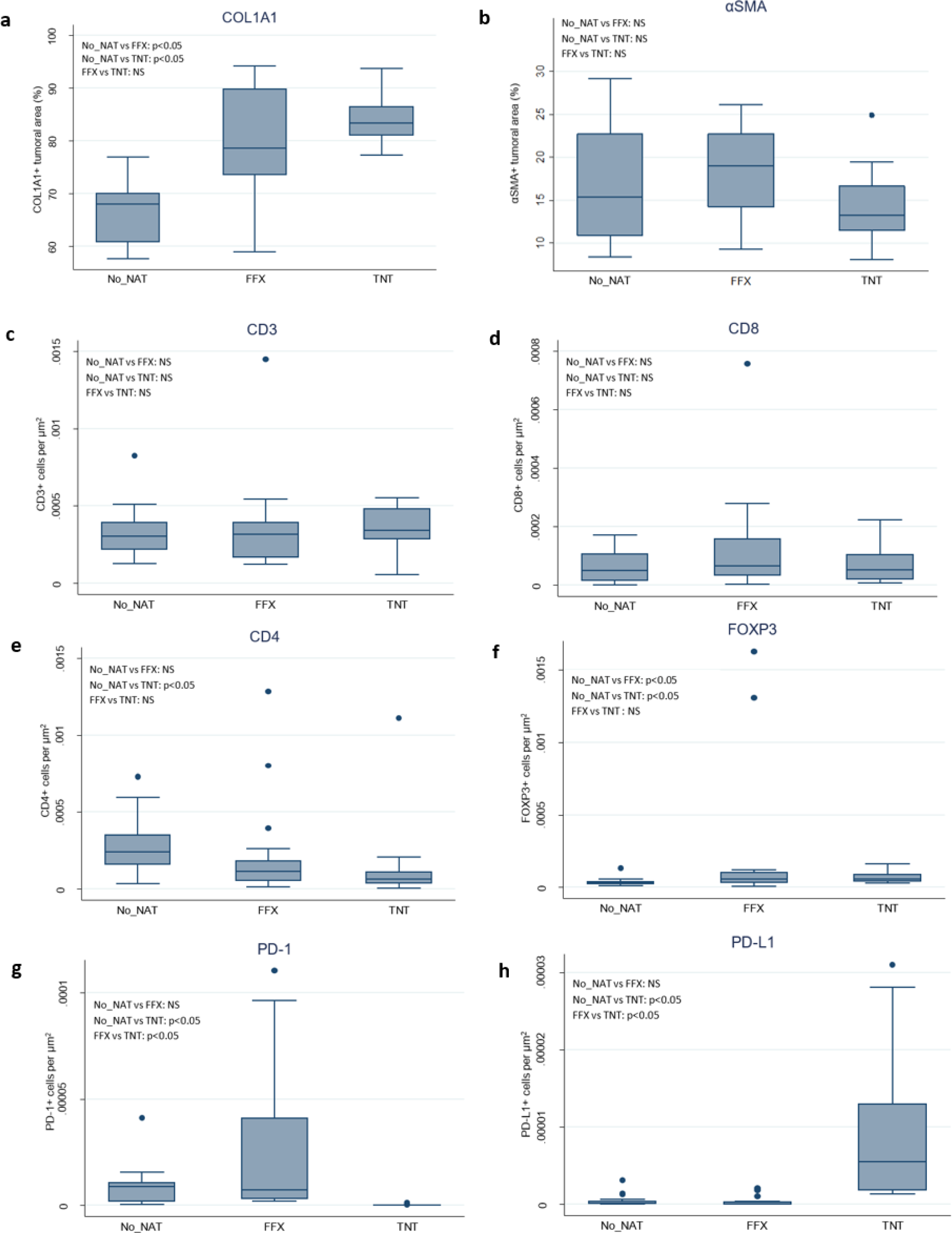
IHC immune and stromal profiling of our whole cohort (n=50). No_NAT: untreated; FFX: FOLFIRINOX; TNT: Total neoadjuvant treatment (FFX + iHD-SBRT); NS: non-significant

**Figure 6.**
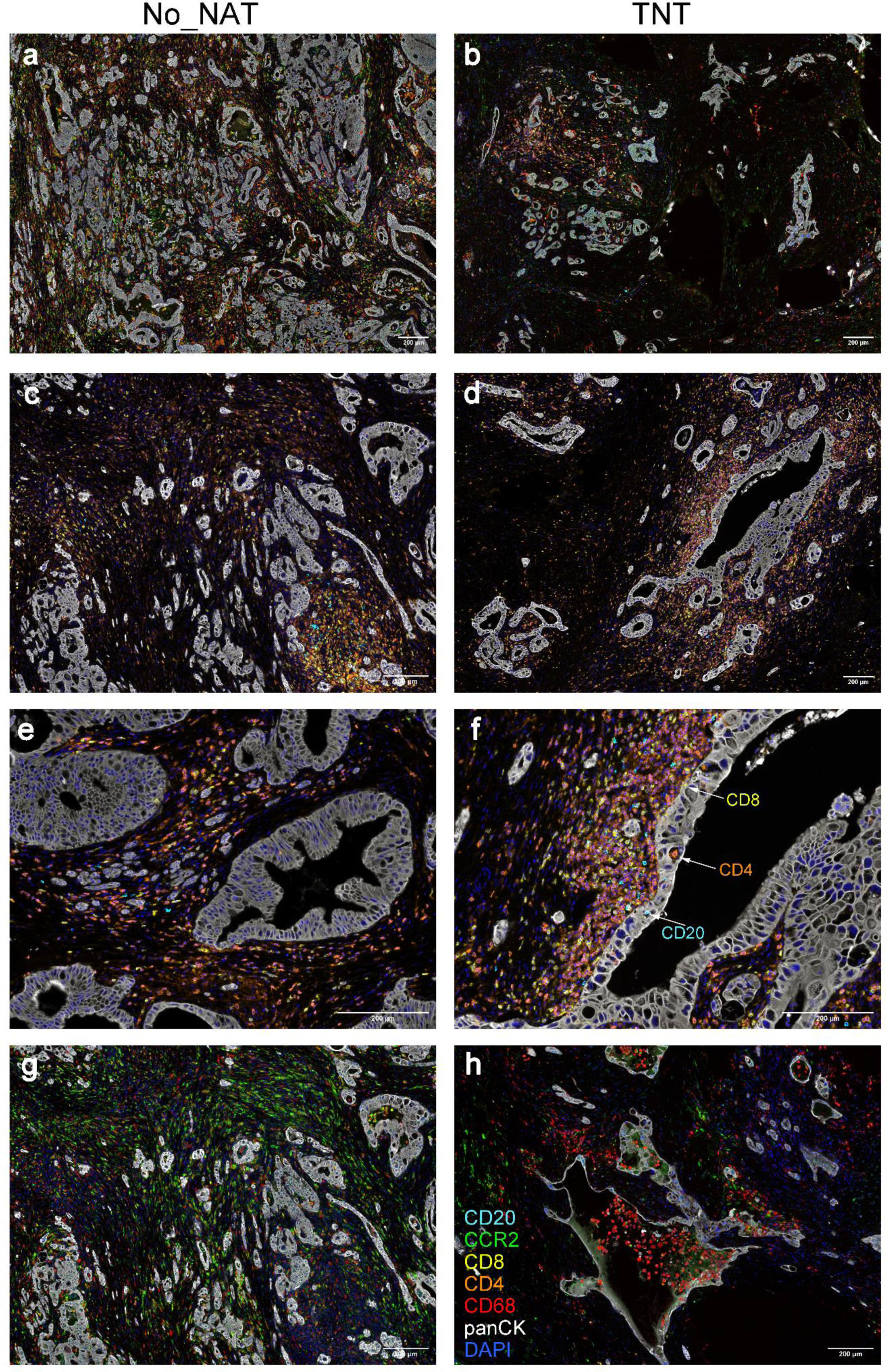
6-plex panel + DAPI multiplex IHC in No_NAT and TNT samples (n= 4). Representative images of: **(b)** Global immune infiltration in No_NAT group with high density of tumoral glands; **(c)** Global immune infiltration in TNT group with less density of tumoral glands; **(d)** Global distribution of tumor infiltrating lymphocytes (TILs) in No_NAT group; **(e)** TILs in TNT group are not sequestrated within the collagenous area; **(f)** TILs in No-NAT group close to the tumoral glands; **(g)** TILs in TNT group are mainly located close and in direct with the tumoral glands; CD4+, CD8+ and CD20+ was observed within the tumoral glands. Cell swelling and pyknotic nucleus of the tumoral cells can be observed in TNT treated PDAC; **(h)** Tumor associated macrophages (TAMs) and CCR2+ cells populations in No_NAT group; **(i)** TAMs are frequently observed within the lumen of tumoral glands in TNT group and CCR2+ cells expression is maintained. No_NAT: untreated; TNT: Total neoadjuvant treatment (FFX + iHD-SBRT)

### Immunosuppressive cells remain present after both neoadjuvant treatments

IHC stainings were performed to explore the immunosuppressive populations of pan-macrophages CD68, CCR2, FOXP3 and PD-1/PD-L1 markers in the TME of our whole PDAC cohort. After different neoadjuvant treatments, no significant differences were observed for CD68+ and CCR2 + cells while the expression of FOXP3 was significantly increased in both TNT and FFX group compared to No_NAT (**Fig. 5f**, **Fig. 6** and **Supplementary Fig. 6**). In the TNT group, CD68+ cells were frequently visualized within the lumen of the remaining tumoral glands (**Fig. 6h**). Expression of PD-L1 and PD-1 was scarce on our whole cohort. PD-L1 expression was significantly increased in the TNT cohort compared to both No_NAT and FFX group but its expression on lymphocytes-like cells remained globally low and weak in the TNT group with a majority of the samples being negative. On the other side, PD-1 expression was significantly decreased and almost null in the TNT group compared to No_NAT and FFX groups. (**Fig. 5g-h** and **Supplementary Fig. 6)**

### xCell deconvolution analysis of the immune TME shows decreased CD4 Th2 population and increased macrophages polarity after TNT

Given the significant decrease in the CD4+ population after TNT demonstrated by IHC data, the presence of the signatures of various T helper cells sub-populations was explored through xCell deconvolution analysis. The results revealed a significant reduction in the CD4 Th2 population in the TNT group compared to the FFX group (**Fig. 7**). In consistence with the IHC data, xCell analysis of the global macrophage population marked no difference among the groups. However, a significant increase in both, M1 and particularly M2-macrophage sub-populations was observed in the TNT samples compared to the FFX group (**Fig. 7**). Conversely, myeloid dendritic cells (MDCs) were significantly decreased after TNT compared to FFX alone, while no significant differences were observed for the neutrophil population (**Fig. 7** and **Supplementary Fig. 7**).

**Figure 7.**
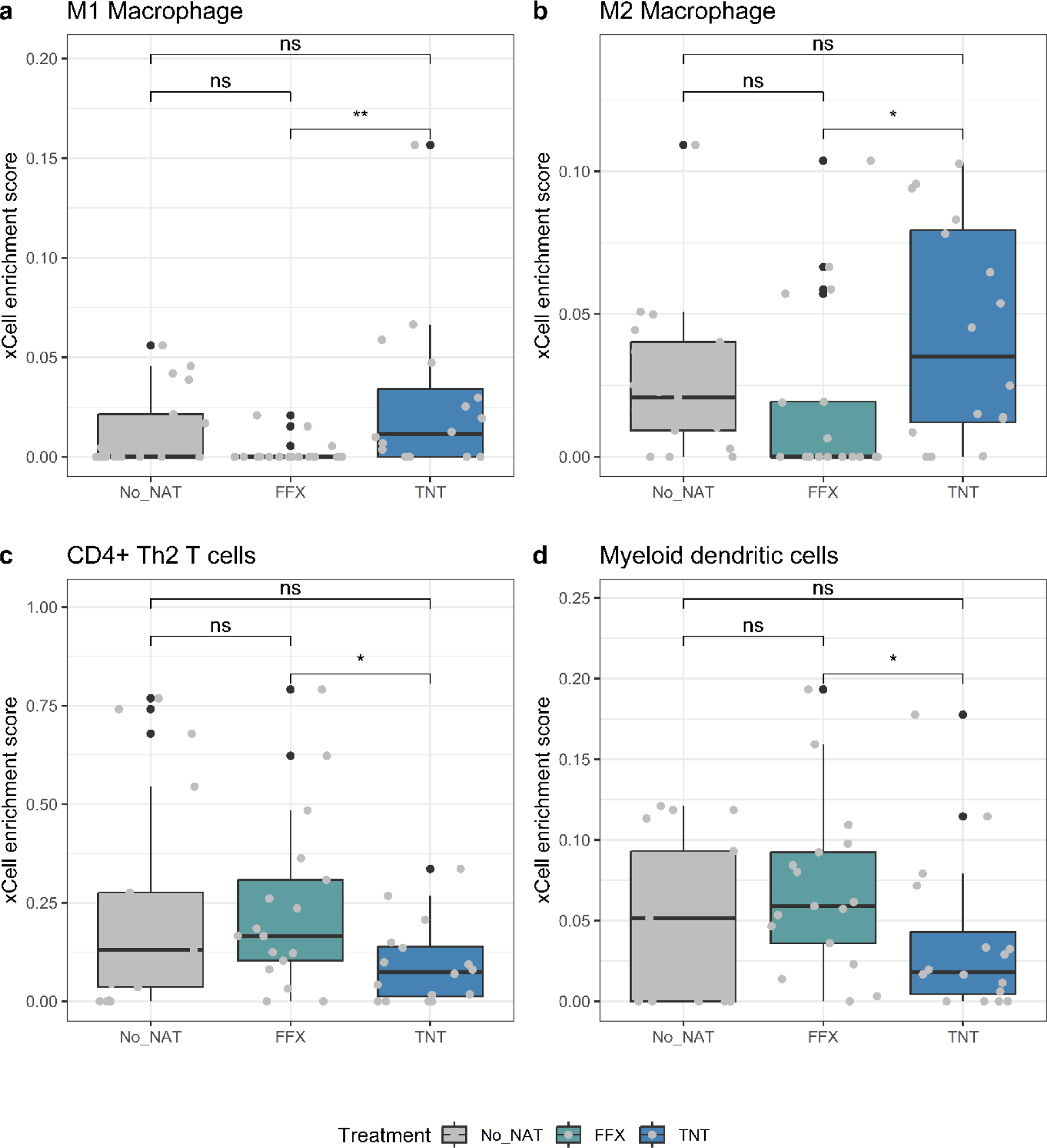
Cell type enrichment analysis using xCell. Cell type enrichment analysis performed using xCell deconvolution showing a significant enrichment of M1-tumor associated macrophages (TAMs) (p=0.0045) **(a)** and M2-TAMs (p=0.024) **(b)** in TNT vs FFX samples. A significantly lower enrichment of CD4+ Th2 T cells (p=0.029) **(c)** and myeloid dendritic cells (p=0.032) **(d)** were observed in the TNT vs FFX group. No_NAT: untreated; FFX: FOLFIRINOX; TNT: Total neoadjuvant treatment (FFX + iHD-SBRT); NS: not significant.

## DISCUSSION

TNT incorporating modern multi-agent chemotherapy, in particular FFX, and innovative radiotherapy such as (nearly) ablative SBRT, has shown promising oncological results in PDAC and is currently investigated in several ongoing prospective randomized trials, including ours. [7, 12–18] Indeed, even in the limited cohort of our study, the 1y-DFS was still statistically in favor of the TNT group (87.5 vs 70.6 vs 41.2% for TNT, FFX and No_NAT, respectively, p=0.017). Despite including significantly more LA patients with larger tumor diameter at diagnosis, the TNT group displayed favorable median DFS and OS. Nonetheless, further well-designed trials, combining these treatments with targeted therapies and stratified treatment approaches, are urgently needed to improve the dismal patients’ prognosis. [5] For this purpose, we hereby investigated for the first time the histo-molecular modulations induced by FFX alone and FFX followed by iHD-SBRT (TNT group).

We identified distinct gene expression patterns and key-pathways, clearly distinguishing two different transcriptional profiles after neoadjuvant treatment with FFX alone or followed by iHD-SBRT. Notably, high-dose SBRT demonstrated the ability to counteract and reverse many of the detrimental transcriptional modulations associated with FFX. While FFX alone led to an increased expression of unfavorable processes linked to EMT, angiogenesis, histone demethylation and intracellular transport of glucose, the addition of iHD-SBRT reversed these effects. Furthermore, metabolic profiles differed based on the neoadjuvant treatment received, with TNT more associated with an increased mitochondrial activity and a more favorable cholesterogenic metabolism compared to FFX alone. [29] These findings provide an additional rationale for combining high-dose SBRT with FFX and may partially explain the promising oncological outcomes obtained with this approach.

In the past decade, transcriptomic-driven subtyping of PDAC was performed by several groups, including ours, using different classification names. [19–25] *In fine*, two main molecular subtypes were systematically identified in these studies: the “Classical” and the “Basal-like” subtype (also denominated as squamous or quasi-mesenchymal). [19–25] The latter is associated with a poorer prognosis, less differentiated tumors and displayed characteristics of EMT. [19–25] On the opposite, the “Classical” subtype is usually associated with better survival outcomes and well-differentiated tumors. [19–25] Although data are still scarce and require further validation in PDAC, the response to therapies seems different according to the molecular subtypes. [24, 26, 28–29] In particular, it is suggested that FFX provides a better response (DFS) in the “Classical” subtype compared to Basal-like subtypes for which gemcitabine-based chemotherapy seems more effective. [19, 23, 26, 31–33] In our study, we observed a significant enrichment in “Basal-like” and active stroma signatures after induction therapy with FFX only. These results are in concordance with the literature, and in particular, with the study by Porter *et al*. that demonstrated in PDAC cell lines a shift from the Classical toward the Basal-like state after FFX treatment. [33–34] To the best of our knowledge, this is a first study investigating potential reprogramming of molecular expression following high-dose SBRT (> 35Gy in 5 fractions) in PDAC. Interestingly, we observed with the addition of iHD-SBRT to FFX a significant enrichment shift toward the “Classical subtype”, related to better prognosis, which was consistent through various signatures available and the molecular gradient PAMG score. [28] To date, the only study exploring molecular subtypes in patients treated with RT is the recent single-nucleus RNAseq study by Hwang *et al.*. This study analyzed 43 PDAC patients; 18 with NT tumors and 25 having received highly variable types of neoadjuvant treatments (including conventional CRT + FFX +/- losartan [n=19] and two patients treated with FFX + low-dose SBRT [33Gy in 5 fractions] + losartan +/- nivolumab). [25] Although non-significant, the authors reported a lower expression of the Squamoid program (similar to “Basal-like”), in the CRT group compared to No_NAT group, supporting our findings. These data also highly suggest that high-dose SBRT targets the “Basal-like” subpopulation more effectively (selection) and/or reprograms the “Basal-like” population induced post-FFX into a more “Classical-like” one (reprogramming). This molecular plasticity process could be mediated through TGFβ activity, as suggested by our transcriptomic data. Indeed, TGFβ has been implicated as a key regulator of cancer cell plasticity between the “Basal” and “Classical” states in PDAC mouse models, with the TGFβ blockade promoting the “Classical” state with increased chemosensitivity. [35]

One of the main transcriptomic modulations observed after neoadjuvant treatments involves stroma remodeling. After iHD-SBRT, compared to both NT and FFX groups, a clear shift towards a more normalized stroma associated with better prognosis was noted. This prompted further investigation into several key-stromal components. Notably, the deposition of ECM, particularly collagen I, significantly increased after neoadjuvant treatments as evidenced by RNAseq / IHC analyses and corroborated by previous studies [36–37]. While an important desmoplasia was previously thought to be only a contributor of tumor progression due to factors such as increased of interstitial fluid pressure, barrier to immune intratumoral infiltration and drug delivery, recent findings suggest that an increased stromal compartment could correlate with a better survival and restrain progression, depending on the cells of its origin. [36–42] In untreated PDAC, the complex and heterogeneous CAF population is the main origin of the desmoplasia (≈90%) but their modulations induced by RT are almost unknown. [41,42] Despite observing a significant increase in COL1A1, the population of myCAFs, reputed to be the subtype most involved in ECM deposition and associated with poor prognosis, was not increased post iHD-SBRT as evidenced by both RNAseq and IHC analysis. [43–45] These data suggest either a simple enhancement of myCAFs activities and/or a potential increase in external collagen production by other cell types. Furthermore, the iCAF subpopulation increased after neoadjuvant treatments, including iHD-SBRT, aligning with recent data from Zhou *et al.* who reported a similar increase in iCAFs in chemotherapy treated samples (n=14; FFX and/or gemcitabine/nab-paclitaxel and 1 case with conventional CRT). [46] High-expression of iCAFs was associated with improved prognosis in other No_NAT PDAC cohorts, which was validated in our study in two independent No_NAT cohorts. [43, 47–48] We further demonstrated a significant association between iCAF-high population and a better DFS after neoadjuvant treatment with TNT, confirming its potential prognostic /predictive role in PDAC. Given that different neoadjuvant treatments generate different effects on the CAF populations, the effectiveness of the addition of therapies targeting CAFs in PDAC may vary depending on the treatment combination used and studies should be encouraged to explore this field.

After iHD-SBRT, the T-lymphocytes infiltration including cytotoxic CD8+ T cells was globally preserved, with immune cells still able to infiltrate close to, and even in direct contact with the tumoral cells despite increased desmoplasia. Previously, Mills *et al*. assessed the CD4/CD8 infiltration within or beyond the areas of dense collagen in a small cohort of nine patients treated with low-dose SBRT only (25Gy in 5 fractions). The authors reported fewer T-cells in these areas in treated samples compared to No_NAT samples, suggesting that T-cell sequestration is not promoted post-SBRT. [36] Another study identified several immune cell marker differences after neoadjuvant treatments, including 12 patients treated with RT, in different area of the tumor through spatial analysis. [49] As expected, we observed an increase in immunosuppressive populations after TNT (notably FOXP3+ Treg cells and macrophages M2-sub-population), however MDCs, PSCs and CD4-Th2 cells were decreased. Finally, the expression of PD-1/PD-L1 was scarce in our whole cohort and, particularly after iHD-SBRT, with almost no expression of PD-1 while PD-L1 increased but remained rare. Consequently, our data do not support the use of anti-PD-1/PD-L1 in PDAC, including in combination with FFX or TNT. Indeed, to date, the association of PD-1/PD-L1 inhibitors with chemotherapy +/- RT remains a failure in PDAC clinical trials. [5–6]

Despite being constrained by several factors, including the absence of matched pre- and post-treatment specimens and limited sample size, our study demonstrates for the first time that high-dose SBRT is capable of durable and in-depth remodeling of PDAC, at the stromal, metabolic and molecular levels. The main significant alterations identified following TNT are resumed in **Fig. 8**, including the capability of reversing several unfavorable enrichment/activations induced by chemotherapy, supporting its complementarity with FFX, along with the potential immune/targeting therapies to be associated with a TNT strategy. This work provides comprehensive insight into human PDAC to more accurately guide the development of new combination strategies involving SBRT. Prospective evaluation of our results will be conducted in the ongoing randomized phase II STEREOPAC trial, planning to enroll 256 patients diagnosed with BR tumors (FFX +/- iHD-SBRT). [16] Further investigation into the exact mechanisms involved in all the reprogramming and alterations induced in PDAC by high-dose SBRT should be pursued in preclinical models and human matched pre- and post-treatment specimens.

**Figure 8.**
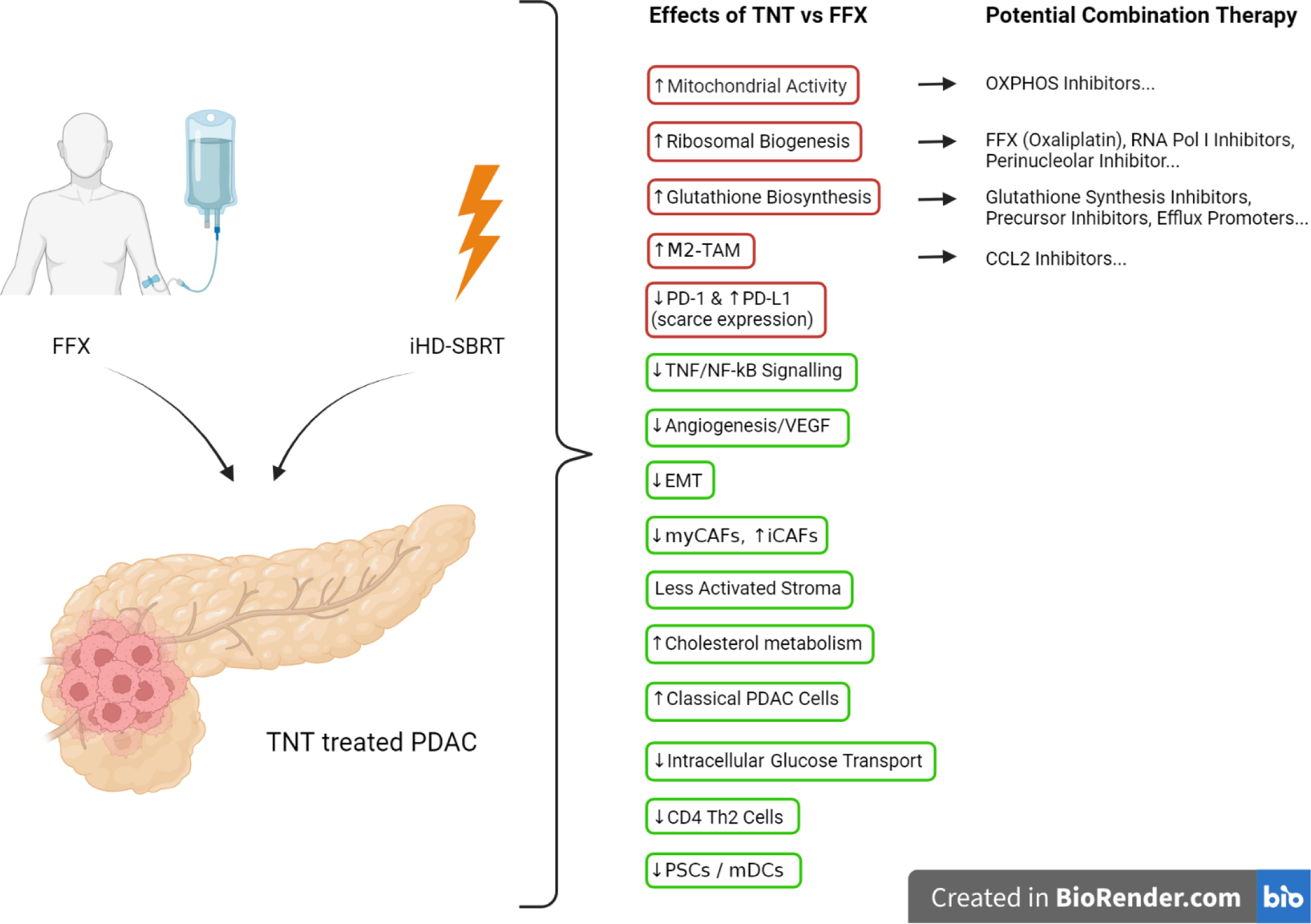
Main identified immuno-molecular modulations following TNT compared to FFX alone in PDAC and selected potential targeted therapy to be combined with TNT. M2-TAM: M2 polarized tumor associated macrophages; EMT: epithelial to mesenchymal transition; myCAF: myofibroblastic cancer associated fibroblast; iCAF: inflammatory cancer associated fibroblast; PDAC: pancreatic ductal adenocarcinoma; PSC: pancreatic stellate cell; MDC: myeloid dendritic cell

## METHODS

### Patients

This study included the use of residual tissue from 50 resected PDAC tumors in Erasme and Pitié Salpêtrière hospitals. All patients had surgery between 2011 and 2020 and archived formalin fixed paraffin-embedded (FFPE) tumor specimens from surgery were available. The main inclusion criteria were patients of age ≥ 18 with complete clinicopathological data available, no evidence of metastatic disease prior to surgery, patients having received no neoadjuvant treatment (No_NAT group), an induction chemotherapy with FFX only (FFX group) or patients treated with a TNT including FFX followed by iHD-SBRT before surgery (TNT group). The main clinical exclusion criteria were the use of any other neoadjuvant treatment (including in case of shift to another type of neoadjuvant chemotherapy such as gemcitabine/nab-paclitaxel), a tumor histology other than a ductal adenocarcinoma (including PDAC associated with intraductal papillary mucinous neoplasm [IPMN]) and patients who died from postoperative complications within 30 days after surgery.

### Data Collection

An aggregated retrospective database with standardized clinicopathological variables was created for patients resected in Erasme and Pitié Salpêtrière hospitals. The variables included: sex, age at diagnosis, level of CA19.9 at diagnosis, clinical disease stage, tumor site, preoperative treatments received, type of surgical resection, TNM classification, histological grade, lymphovascular and perineural invasion, and relevant outcomes parameters.

### Neoadjuvant treatment

Patients receiving a neoadjuvant treatment included an induction with FFX chemotherapy regimen for a median of 6 cycles. The FFX regimen consisted in an intravenous infusion of oxaliplatin (85mg/m^2^, 2h) then an intravenous infusion of leucovorin (400mg/m^2^, 2h) concomitantly with a 90-min intravenous infusion of irinotecan (165-180mg/m^2^) followed by a 46h continuous infusion of fluorouracil (2000-2400mg/m^2^), and was given once every two weeks.

For sixteen patients, FFX was followed by iHD-SBRT as previously described in details in [14, 50], according to the TNT strategy implemented in our hospital since January 2018 for localized PDAC. A surgical exploration was performed in case of no progression 4 to 7 weeks after iHD-SBRT. Briefly, the SBRT treatment was designed to individually maximize the dose prescribed to the tumor and vessels interfaces (D_max(0.5cc)_ <53Gy in 5 fractions) while following an isotoxic dose prescription (IDP). In an IDP, the dose prescription is not based on the coverage of the planning target volume (PTV) but on the predetermined limiting dose constraints to the neighbouring organs at risks in order to control toxicity. [14, 50] The following dose constraints were applied: for planning organ at risk volumes (PRVs) stomach, duodenum, colon and small bowel, D_max (0.5cc)_ <35Gy, V_30Gy_<2cc; PRV spinal cord, V_20Gy_<1cc and for kidneys, D_mean_<10Gy and V_12Gy_<25%).

### Sample processing and RNA isolation

For accurate reference slides, new FFPE tissue section was cut at 4µm then stained with H&E for all the representative tumoral blocks identified by specialized gastrointestinal pathologists (LV, PDM, NH). Tissue sections were scanned using a Nanozoomer 2.0-RS Digital slide scanner (Hamamatsu). The H&E digital slides used as reference were reviewed by CB and a specialized pathologist (LV) to delineate the tumoral area prior to RNA isolation. From the 50 FFPE blocks, five consecutive 6-8µm non-stained slides were cut in RNAse free conditions. The tumoral area was then demarcated on each slide, directly comparing it with the reference H&E slide.

The delineated tumoral sections were manually scrapped and RNA was extracted from the scrapped sections with the ALLPrep FFPE tissue kit^©^ following the manufacturer’s instructions for semi-automated RNA extraction via Qiacube instrument (Qiagen, Venlo, The Netherlands). RNA samples were run on an Agilent 2100 bioanalyzer using the RNA 6000 Pico LabChip kit (Agilent, Diegem, Belgium). The bioanalyzer electropherograms were analyzed by Agilent 2100 Expert Software to determine the RNA quantity and quality. RNA samples with DV200 >30% were selected and 100 ng of RNA was used for the library preparation. NGS libraries were prepared using the QuantSeq Library Prep Kit for Illumina (Lexogen) as per manufacturer recommendations’. The libraries were sequenced on NovaSeq using NovaSeq 6000 S2 Reagent Kit with 100 bp single reads.

### RNA-sequencing Data Analysis

FASTQ files were checked for sequencing quality via FastQC. [51] The quantification of transcript abundance was done from the raw RNA-seq files using the Kallisto v0.50.0 pseudo-alignment method. [52] Kallisto was performed with a 100-bootstrap value, using a transcriptome index constructed from the human reference transcriptome GRCH38 from Ensembl. Gene-level quantification of estimated counts was performed using the R-package tximport v1.26.1. (data available here: <u>10.5281/zenodo.10939866</u>) [53] Poorly covered genes (read count <10 in more than half of the samples) were removed for further analysis. Differential gene expression (DGE) analyses were performed between patients that received different treatments using the R-packages edgeR v3.40.2 and limma v3.54.2 packages. [54–55] Heatmap representations of the genes with a p-value lower than 0.05 in ech of the comparisons applied in the DGE analyses were generated using Complex Heatmap v2.14.0 package (**Supplementary Fig. 8**). [56] The PAMG classifier was applied to determine the chemosensitivity and the agressivity of the samples. [28]

### Functional analysis

With the aim of characterizing the molecular characteristics of each neoadjuvant therapy, Gene Set Enrichment Analysis (GSEA) was performed on a pre-ranked list of genes using the fgsea R package v1.24.0. [57] Only enrichments of gene sets with a padj< 0.05 were considered as significant. Gene signatures of PDAC and cancer associated fibroblast (CAF) subtypes were collected from the CancerRNASig package. Molecular Signature Database (MSigDB), Ontology and Canonical pathways gene sets were obtained by the msigdb package v1.6.0. Gene Set Variation Analysis (GSVA) was applied to samples as a single sample classifier of different CAF subtypes. [58] Finally, immune cell fractions were estimated by the xCell algorithm and statistical analysis between treatments of the immune populations was obtained by the package ggpubr v0.6.0. [59–60]

### Immunohistochemistry (IHC)

FFPE full-face tissue sections (4µm) from the 50 tumors were single and dual immunohistochemically stained for CD3/CD20, CD4/CD8, PD-1/PD-L1, CD68, CCR2, FOXP3, COL1A1, and αSMA. All antibodies and their dilution are listed in the **Supplementary Table 1**. Chromogenic IHC (cIHC) were performed on a Ventana Benchmark XT automated staining instrument with the ultraVIEW DAB and ultraVIEW Red Detection Kit (Ventana Medical Systems). All antibodies were initially tested on positive and negative control tissues and staining patterns were validated by pathologists (LV and PDM). cIHC slides were acquired at 40x with a Nanozoomer 2.0-RS Digital slide scanner (Hamamatsu). Delineation of the tumor area was performed by CB and verified by two experimented specialized pathologists (LV and PDM). Quantification of the different stainings was performed with the Visiopharm^©^ software.

### Multiplex Immunohistochemistry (mIHC)

FFPE tissue sections (4µm) were processed manually for mIHC using Opal reagents (Akoya Biosciences) for illustration purposes in four representative samples (2 in the No-NAT and TNT groups). Briefly, slides were first heated at 37°C overnight before being deparaffinized hydrated through an ethanol gradient and fixed in 10% neutral buffer formalin. Heat-induced antigen retrieval was achieved in Antigen Retrieval (AR) 9 buffer (Akoya Biosciences) using a microwave (Panasonic with Inverter technology). Slides were labeled for CD20 (B cells), CD4 (Th cells), CD8 (cytotoxic T cells), CD68 (macrophages), CCR2 (chemokine CCL2 receptor), pan-cytokeratin (cancer cells) and DAPI (all nuclei) according to the manufacturer’s instructions (Opal 6-Plex Manual Detection Kit - for Whole Slide Imaging, NEL861001KT, Akoya Biosciences) (**Supplementary Table 1**). Slides were mounted with Prolong Diamond Antifade Mountant (Life Technologies Europe BV). The whole slides were acquired with the PhenoImager HT scanner (Akoya Biosciences) using appropriated exposure times. Tonsil tissue was used as positive control. Region of interests (ROIs) were selected in PhenoChart Whole Slide Viewer by an experimented gastrointestinal pathologist (LV). ROIs were unmixed using the synthetic spectral library and the tissue autofluorescence extracted from an unstained PDAC was removed in inForm Tissue Analysis Software (V.2.6.0, Akoya Biosciences).

### Statistical analysis

Statistical analyses were performed using Stata 14 and R. Data normality was assessed using histograms, boxplots, and quantile–quantile plots, and the equality of variances was checked using the Levene’s test.

Categorical data were presented as percentages and numbers, while continuous data were described using median and P25–P75, and due to asymmetric distribution, analyzed with nonparametric tests such as the Kruskal-Wallis rank test for group differences. Chi² tests were employed for categorical data. Bonferroni corrections were applied following multiple comparisons between the different groups.

Survival analyses were conducted using the survival v3.5-3 and survminer v0.4.9 packages. Log-rank test was used to calculate the differences in Kaplan-Meier curves and p-values < 0.05 were considered as statistically significant. Multivariate Cox proportional hazard regression models were applied for survival with a 95% confidence interval. OS was defined as the time in months from diagnosis to death due to cancer recurrence. DFS was defined as the time from diagnosis to the first documentation of recurrent disease following surgery. Loco-regional DFS (LR-DFS) was defined as the time from diagnosis to the first documentation of loco-regional recurrence (in the original tumor location or the N1-2 lymph node areas).

Non-parametric Wilcoxon test in R v4.2.3 and RStudio v2023.3.0.386 environments was used for RNAseq data analysis, assessing significant differences in treatments in PAMG, Puleo components projections, and xCell immune deconvolution outputs with p values < 0.05 considered statistically significant.

### Study approval

This study was approved by the Institutional Review Board of Erasme University Hospital and Pitié Salpêtrière hospital under the approval numbers P2018/392 - A2020-115 and 2014/58NICB respectively.

## Acknowledgements

The first authors disclosed receipt of the following financial support for the research: this work is supported by doctoral grants from the “Les Amis de l’Institut Bordet / L’Association Jules Bordet” grant numbers: [2019–31] (CB, DVG, LM) and [2021–03] (CB), by the “Fonds de la Recherche Scientifique – FNRS” grant numbers [FC 33593] (CB) and [PDR T0011.22] (OAS). JN was financially supported by Fonds Erasme and VD by a grant of the “Fondation Contre le Cancer - FCC” grant number [F/2020/1402]. IR and JLVL supported this work through a FCC grant number [FAF-C/2018/1203].

## Author contributions

This study was designed and conceptualized by CB, TA and JLVL. JN, LV, PDM, ND and JBB provided human samples. Clinical data were collected by CB, JN, SZ and LMa. (m)IHC experiments, quantification, data analysis, interpretation and related figure design were done by CB, LV, AB, MH, KS, IR, PDM and KWG. RNA isolation, RNAseq data analysis, interpretation and related figures were done by CB, OAS, JN, EQ, VT, VD, RN and TA. CB, TA and OAS drafted the manuscript. CB, DVG, LMo, IR and JLVL obtained funding for the study. Editing was performed by LMo and DVG. All authors performed critical revisions. All authors read and approved the final manuscript.

