## Supplementary Data 1 for "Favorable histo-molecular remodeling of pancreatic ductal adenocarcinoma after Total Neoadjuvant Therapy including Stereotactic Body Radiotherapy"

| pathway | pval | padj | log2err | ES | NES | size |
|---|---|---|---|---|---|---|
| CAF_FMG20_CAF.S1 | 0,001372 | 0,005647 | 0,45506 | -0,4259 | -1,7732 | 72 |
| CAF_FMG20_IFNγ.iCAF | 0,265849 | 0,447999 | 0,11829 | 0,39425 | 1,16686 | 16 |
| CAF_FMG20_IL.iCAF | 0,468451 | 0,657633 | 0,0802 | -0,2963 | -0,9901 | 25 |
| CAF_FMG20_Normal.Fibroblast | 0,071269 | 0,160082 | 0,25296 | 0,2982 | 1,30843 | 80 |
| CAF_FMG20_TGFβ.myCAF | 0,031334 | 0,081693 | 0,32178 | -0,4738 | -1,5834 | 25 |
| CAF_FMG20_detox.iCAF | 0,973025 | 0,986539 | 0,04522 | -0,1853 | -0,5917 | 21 |
| CAF_FMG20_ecm.myCAF | 0,000631 | 0,003073 | 0,47727 | -0,5372 | -1,937 | 36 |
| CAF_FMG20_wound.myCAF | 0,005552 | 0,019299 | 0,40702 | -0,5073 | -1,7725 | 31 |
| CAF_Neuzillet22_MYH11 | 0,922631 | 0,98637 | 0,04783 | -0,2019 | -0,6961 | 29 |
| CAF_Neuzillet22_PDPN | 0,521154 | 0,704523 | 0,07493 | -0,2945 | -0,948 | 22 |
| CAF_Neuzillet22_POSTN | 0,260504 | 0,447999 | 0,12154 | 0,32723 | 1,13225 | 28 |
| CAF_Neuzillet22_subPOSTN | 0,98167 | 0,98844 | 0,04735 | 0,19248 | 0,59906 | 19 |
| CAF_Turley20_hCAF0TGFB | 3,74E-08 | 6,07E-07 | 0,71951 | -0,4634 | -2,1414 | 134 |
| CAF_Turley20_hCAF1early | 0,006417 | 0,021789 | 0,40702 | -0,3217 | -1,5 | 148 |
| CAF_Turley20_hCAF2il6lif | 0,423701 | 0,618604 | 0,07687 | -0,1996 | -1,0099 | 258 |
| CAF_Tuveson19_iCAF | 0,613971 | 0,765259 | 0,0645 | -0,2172 | -0,9165 | 76 |
| CAF_Tuveson19_myo | 0,1537 | 0,297121 | 0,15419 | -0,4455 | -1,302 | 15 |
| CCCA_MetaProg_MP1.Cell.Cycle.G2.M | 0,414013 | 0,610564 | 0,09285 | 0,27942 | 1,03026 | 36 |
| CCCA_MetaProg_MP10.Protein.maturation | 0,026411 | 0,071406 | 0,35249 | 0,41744 | 1,55701 | 38 |
| CCCA_MetaProg_MP11.Translation.initiation | 0,276956 | 0,447999 | 0,11777 | 0,2998 | 1,11824 | 38 |
| CCCA_MetaProg_MP12.EMT.I | 1,07E-05 | 8,68E-05 | 0,59333 | -0,5624 | -2,1651 | 47 |
| CCCA_MetaProg_MP13.EMT.II | 0,085502 | 0,183578 | 0,20896 | -0,353 | -1,3357 | 44 |
| CCCA_MetaProg_MP14.EMT.III | 0,447598 | 0,643009 | 0,08999 | 0,25916 | 1,00703 | 45 |
| CCCA_MetaProg_MP15.EMT.IV | 0,103774 | 0,216442 | 0,19002 | -0,3528 | -1,3112 | 42 |
| CCCA_MetaProg_MP16.MES.glioma. | 0,001393 | 0,005647 | 0,45506 | -0,491 | -1,8248 | 42 |
| CCCA_MetaProg_MP17.Interferon.MHC.II.I. | 0,27907 | 0,447999 | 0,11725 | 0,29739 | 1,11631 | 39 |
| CCCA_MetaProg_MP18.Interferon.MHC.II.II. | 0,603814 | 0,759972 | 0,07236 | 0,2428 | 0,91785 | 41 |
| CCCA_MetaProg_MP19.Epithelial.Senescence | 0,080851 | 0,178852 | 0,23113 | 0,35216 | 1,32937 | 40 |
| CCCA_MetaProg_MP2.Cell.Cycle.G1.S | 0,199575 | 0,368835 | 0,14206 | 0,32276 | 1,19005 | 36 |
| CCCA_MetaProg_MP20.MYC | 0,115217 | 0,236926 | 0,19381 | 0,32135 | 1,26728 | 48 |
| CCCA_MetaProg_MP21.Respiration | 0,00067 | 0,003157 | 0,47727 | 0,5016 | 1,88288 | 39 |
| CCCA_MetaProg_MP22.Secreted.I | 0,366667 | 0,563509 | 0,10282 | 0,26973 | 1,0528 | 46 |
| CCCA_MetaProg_MP23.Secreted.II | 0,059158 | 0,137096 | 0,32178 | 0,36283 | 1,43083 | 48 |
| CCCA_MetaProg_MP24.Cilia | 0,817227 | 0,938441 | 0,05725 | 0,22567 | 0,78084 | 28 |
| CCCA_MetaProg_MP25.Astrocytes | 0,594142 | 0,759972 | 0,07254 | 0,25193 | 0,91336 | 33 |
| CCCA_MetaProg_MP26.NPC.Glioma | 0,972632 | 0,986539 | 0,04929 | 0,16959 | 0,61147 | 32 |
| CCCA_MetaProg_MP27.Oligo.Progenitor | 0,655462 | 0,778028 | 0,06783 | 0,25708 | 0,88954 | 28 |
| CCCA_MetaProg_MP28.Oligo.normal | 0,783366 | 0,909762 | 0,05536 | -0,2337 | -0,8164 | 31 |
| CCCA_MetaProg_MP29.NPC.OPC | 0,307547 | 0,488064 | 0,10358 | -0,2929 | -1,0862 | 41 |
| CCCA_MetaProg_MP3.Cell.Cylce.HMG.rich | 0,408207 | 0,609462 | 0,09466 | 0,26966 | 1,03076 | 43 |
| CCCA_MetaProg_MP30.PDAC.classical | 0,002561 | 0,009589 | 0,43171 | 0,46127 | 1,77893 | 44 |
| CCCA_MetaProg_MP31.Alveolar | 0,030189 | 0,080138 | 0,35249 | 0,42837 | 1,52268 | 30 |
| CCCA_MetaProg_MP32.Skin.pigmentation | 0,856842 | 0,954954 | 0,05514 | 0,20832 | 0,75111 | 32 |
| CCCA_MetaProg_MP33.RBCs | 0,650628 | 0,778028 | 0,06799 | 0,24415 | 0,88514 | 33 |
| CCCA_MetaProg_MP34.Platelet.activation | 0,049336 | 0,118083 | 0,28201 | -0,4153 | -1,4647 | 32 |
| CCCA_MetaProg_MP35.Hemato.related.I | 0,552577 | 0,733421 | 0,0755 | 0,26905 | 0,94161 | 29 |
| CCCA_MetaProg_MP36.IG | 0,35546 | 0,552098 | 0,10245 | 0,30802 | 1,05408 | 27 |
| CCCA_MetaProg_MP37.Hemato.related.II | 0,646729 | 0,778028 | 0,0628 | -0,2485 | -0,8915 | 35 |
| CCCA_MetaProg_MP38.Glutathione | 0,202105 | 0,368842 | 0,14041 | 0,35254 | 1,1952 | 26 |

| | | | | | | |
|---|---|---|---|---|---|---|
| CCCA_MetaProg_MP39.Metal.response | 0,963983 | 0,986539 | 0,04999 | 0,17822 | 0,67371 | 41 |
| CCCA_MetaProg_MP4.Chromatin | 0,9 | 0,980597 | 0,04783 | -0,1997 | -0,7423 | 42 |
| CCCA_MetaProg_MP40.PDAC.related | 0,50431 | 0,688124 | 0,08266 | 0,25223 | 0,97274 | 44 |
| CCCA_MetaProg_MP41.Unassigned | 0,488223 | 0,678862 | 0,08407 | 0,2655 | 0,97147 | 35 |
| CCCA_MetaProg_MP5.Stress | 0,537778 | 0,720326 | 0,08086 | 0,24503 | 0,95642 | 46 |
| CCCA_MetaProg_MP6.Hypoxia | 0,271552 | 0,447999 | 0,12043 | 0,29052 | 1,12043 | 44 |
| CCCA_MetaProg_MP7.Stress.in.vitro. | 0,152577 | 0,297121 | 0,16198 | 0,35685 | 1,24889 | 29 |
| CCCA_MetaProg_MP8.Proteasomal.degradati | 0,043195 | 0,108733 | 0,32178 | 0,37435 | 1,4618 | 47 |
| CCCA_MetaProg_MP9.Unfolded.protein.respo | 0,000257 | 0,001566 | 0,49849 | 0,50939 | 1,94715 | 43 |
| CCK_STIM_Desert-like | 0,036499 | 0,093488 | 0,32178 | 0,37958 | 1,48223 | 47 |
| CCK_STIM_Hepati_Stem-like | 0,988839 | 0,988839 | 0,05122 | 0,15541 | 0,63561 | 60 |
| CCK_STIM_Immune_Classical | 0,630112 | 0,773078 | 0,06378 | -0,2366 | -0,8953 | 44 |
| CCK_STIM_Inflammatory_Stroma | 0,000278 | 0,001612 | 0,49849 | -0,4573 | -1,8796 | 66 |
| CCK_STIM_Tumor_Classical | 0,966728 | 0,986539 | 0,04363 | -0,1691 | -0,6876 | 64 |
| CCK_Sia13_INFLAMGenes | 0,409091 | 0,609462 | 0,0848 | -0,2437 | -1,0321 | 78 |
| CCK_Sia13_PROLIFGenes | 0,010406 | 0,032324 | 0,38073 | -0,2112 | -1,205 | #### |
| DrugBank_Fluorouracil | 0,37395 | 0,568715 | 0,09821 | 0,40115 | 1,08389 | 12 |
| DrugBank_Gemcitabine | 0,156576 | 0,297121 | 0,1608 | 0,50223 | 1,28913 | 10 |
| DrugBank_Irinotecan | 0,098121 | 0,207619 | 0,20659 | 0,54405 | 1,39646 | 10 |
| DrugBank_Oxaliplatin | 0,007431 | 0,024656 | 0,40702 | 0,70681 | 1,75147 | 9 |
| DrugBank_Paclitaxel | 0,918864 | 0,98637 | 0,05019 | 0,23688 | 0,62959 | 11 |
| ECM_Helms22_PSCcaf | 5,63E-06 | 5,14E-05 | 0,61053 | -0,4698 | -2,0294 | 90 |
| IMMU_GenJCI121924_GeneralImmu | 0,785137 | 0,909762 | 0,04698 | -0,1692 | -0,9126 | 482 |
| IMMU_GenJCI121924_ICKrelated | 0,885122 | 0,971638 | 0,04851 | -0,2338 | -0,6698 | 14 |
| IMMU_MCPcounter_B.lineage | 0,967681 | 0,986539 | 0,04486 | -0,2199 | -0,5099 | 7 |
| IMMU_MCPcounter_CD8Tcells | 0,941176 | 0,986539 | 0,0507 | 0,52535 | 0,69844 | 1 |
| IMMU_MCPcounter_Cytotox.lymph | 0,275304 | 0,447999 | 0,11524 | -0,6772 | -1,1618 | 3 |
| IMMU_MCPcounter_Endothelial | 0,211066 | 0,380439 | 0,135 | 0,37921 | 1,19215 | 20 |
| IMMU_MCPcounter_Fibroblasts | 0,024646 | 0,067894 | 0,35249 | -0,6935 | -1,6761 | 8 |
| IMMU_MCPcounter_Mono.lineage | 0,588235 | 0,759972 | 0,07326 | 0,38313 | 0,88372 | 7 |
| IMMU_MCPcounter_Myeloid.dendritic | 0,775665 | 0,909762 | 0,05502 | -0,6129 | -0,819 | 1 |
| IMMU_MCPcounter_NK | 0,640756 | 0,778028 | 0,06896 | 0,68343 | 0,90861 | 1 |
| IMMU_MCPcounter_Neutrophils | 0,618497 | 0,765259 | 0,06644 | -0,3593 | -0,8684 | 8 |
| IMMU_MCPcounter_Tcells | 0,1222 | 0,247794 | 0,18138 | -0,6851 | -1,3787 | 5 |
| PDAC_Bailey16_ADEX | 0,009852 | 0,031269 | 0,38073 | 0,32979 | 1,50273 | 98 |
| PDAC_Bailey16_Immunogenic | 0,934664 | 0,986539 | 0,04433 | -0,172 | -0,7251 | 74 |
| PDAC_Bailey16_Progenitor | 0,156701 | 0,297121 | 0,15965 | 0,35454 | 1,2408 | 29 |
| PDAC_Bailey16_Squamous | 0,000396 | 0,002062 | 0,49849 | -0,2753 | -1,476 | 449 |
| PDAC_CSY20_BasallikeA | 0,847176 | 0,951444 | 0,04477 | -0,1727 | -0,8669 | 245 |
| PDAC_CSY20_BasallikeB | 0,150877 | 0,297121 | 0,14921 | -0,2687 | -1,1892 | 99 |
| PDAC_CSY20_ClassicalA | 0,000694 | 0,003168 | 0,47727 | 0,3025 | 1,5458 | 218 |
| PDAC_CSY20_ClassicalB | 0,191344 | 0,358157 | 0,15115 | 0,25358 | 1,1502 | 96 |
| PDAC_CSY20_Sig11 | 0,864583 | 0,956282 | 0,04586 | -0,1804 | -0,817 | 112 |
| PDAC_CSY20_Sig12 | 0,311072 | 0,48835 | 0,09855 | -0,2364 | -1,073 | 118 |
| PDAC_CSY20_Sig3 | 2,05E-09 | 4,27E-08 | 0,77494 | -0,3348 | -1,8111 | 497 |
| PDAC_CSY20_Sig4 | 0,50116 | 0,688124 | 0,08705 | 0,2072 | 0,98592 | 140 |
| PDAC_CSY20_Sig5 | 0,018846 | 0,053951 | 0,35249 | 0,22137 | 1,23819 | 515 |
| PDAC_CSY20_Sig7 | 0,016943 | 0,049472 | 0,35249 | 0,26295 | 1,33772 | 213 |
| PDAC_CSY20_Sig8 | 0,925566 | 0,98637 | 0,03952 | -0,1631 | -0,8256 | 255 |
| PDAC_CSY20_Sig9 | 0,082979 | 0,180819 | 0,22799 | 0,35162 | 1,3273 | 40 |

| | | | | | | |
|---|---|---|---|---|---|---|
| PDAC_Hwang22_Fibro.Adhesive | 5,66E-09 | 1,03E-07 | 0,76146 | -0,432 | -2,0744 | 180 |
| PDAC_Hwang22_Fibro.Immunomodulatory | 0,831325 | 0,94088 | 0,04726 | -0,1797 | -0,8435 | 153 |
| PDAC_Hwang22_Fibro.Myofibroblastic | 2,19E-13 | 8,01E-12 | 0,94363 | -0,4975 | -2,3922 | 176 |
| PDAC_Hwang22_Fibro.Neurotropic | 0,002143 | 0,008456 | 0,43171 | -0,3164 | -1,5121 | 169 |
| PDAC_Hwang22_Malignlineage.Acinarlike | 4,34E-08 | 6,34E-07 | 0,71951 | 0,41354 | 2,01257 | 162 |
| PDAC_Hwang22_Malignlineage.Basaloid | 4E-07 | 4,87E-06 | 0,67496 | 0,40805 | 1,99945 | 166 |
| PDAC_Hwang22_Malignlineage.Classicallike | 0,463235 | 0,656625 | 0,09466 | 0,20336 | 1,00077 | 173 |
| PDAC_Hwang22_Malignlineage.Mesenchymal | 0,001241 | 0,005328 | 0,45506 | -0,3255 | -1,5558 | 178 |
| PDAC_Hwang22_Malignlineage.Neurallikeprog | 0,449225 | 0,643009 | 0,07687 | -0,2136 | -1,0027 | 153 |
| PDAC_Hwang22_Malignlineage.Neuroendocri | 0,579186 | 0,75501 | 0,07788 | 0,20697 | 0,94311 | 98 |
| PDAC_Hwang22_Malignlineage.Squamoid | 0,000287 | 0,001612 | 0,49849 | 0,34028 | 1,65423 | 158 |
| PDAC_Hwang22_Malignstate.Adhesive | 0,267918 | 0,447999 | 0,10592 | -0,2298 | -1,0955 | 167 |
| PDAC_Hwang22_Malignstate.CyclingG2M | 0,279232 | 0,447999 | 0,10473 | -0,2361 | -1,0919 | 136 |
| PDAC_Hwang22_Malignstate.CyclingS | 0,600707 | 0,759972 | 0,06364 | -0,2045 | -0,9451 | 141 |
| PDAC_Hwang22_Malignstate.Interferonsignali | 0,050347 | 0,11856 | 0,26635 | -0,2721 | -1,274 | 152 |
| PDAC_Hwang22_Malignstate.MYCsignaling | 0,044293 | 0,109606 | 0,28201 | -0,2722 | -1,3014 | 172 |
| PDAC_Hwang22_Malignstate.Ribosomal | 7,08E-14 | 3,45E-12 | 0,95454 | 0,49143 | 2,42956 | 182 |
| PDAC_Hwang22_Malignstate.TNFNFkBsignalin | 6,08E-07 | 6,83E-06 | 0,65944 | -0,4025 | -1,9237 | 169 |
| PDAC_Moffitt15_ActivatedStroma | 7,14E-20 | 5,22E-18 | 1,15122 | -0,3885 | -2,1426 | 646 |
| PDAC_Moffitt15_BasalLike | 7,05E-06 | 6,06E-05 | 0,61053 | -0,2959 | -1,5957 | 469 |
| PDAC_Moffitt15_CellCycle | 0,000587 | 0,002958 | 0,47727 | -0,2112 | -1,2499 | #### |
| PDAC_Moffitt15_Classical | 0,008848 | 0,028706 | 0,38073 | 0,2228 | 1,27193 | 618 |
| PDAC_Moffitt15_Endocrine | 0,000128 | 0,000852 | 0,51885 | 0,27246 | 1,4979 | 397 |
| PDAC_Moffitt15_Exocrine | 0,565327 | 0,743583 | 0,0848 | 0,17707 | 0,96887 | 393 |
| PDAC_Moffitt15_F10 | 0,216718 | 0,384048 | 0,16693 | 0,17784 | 1,04771 | #### |
| PDAC_Moffitt15_F14 | 6,54E-05 | 0,000455 | 0,53843 | 0,22499 | 1,33016 | #### |
| PDAC_Moffitt15_F2 | 8,99E-13 | 2,63E-11 | 0,92143 | 0,29341 | 1,71504 | 981 |
| PDAC_Moffitt15_Immune | 0,000374 | 0,002022 | 0,49849 | -0,255 | -1,4014 | 627 |
| PDAC_Moffitt15_Liver | 0,07 | 0,159688 | 0,27129 | 0,21078 | 1,15927 | 409 |
| PDAC_Moffitt15_Lung | 0,822742 | 0,938441 | 0,04642 | -0,174 | -0,8683 | 233 |
| PDAC_Moffitt15_Muscle | 0,218329 | 0,384048 | 0,15419 | 0,18764 | 1,05981 | 582 |
| PDAC_Moffitt15_NormalStroma | 0,011818 | 0,035947 | 0,38073 | -0,2261 | -1,2504 | 673 |
| PDAC_PDAssigner_Classical | 0,004572 | 0,01628 | 0,40702 | 0,55233 | 1,79206 | 22 |
| PDAC_PDAssigner_Exocrine | 0,002435 | 0,009357 | 0,43171 | 0,59184 | 1,8606 | 20 |
| PDAC_PDAssigner_QM | 0,023988 | 0,067352 | 0,35249 | -0,5604 | -1,638 | 15 |
| PDAC_PDXph1_basal | 4,6E-06 | 4,48E-05 | 0,61053 | -0,2749 | -1,519 | 663 |
| PDAC_PDXph1_classical | 0,0002 | 0,001269 | 0,51885 | 0,26231 | 1,45713 | 477 |
| PDAC_Puleo_Basal | 4,43E-11 | 1,08E-09 | 0,85134 | -0,3283 | -1,8044 | 618 |
| PDAC_Puleo_Classic | 3,01E-05 | 0,000231 | 0,57561 | 0,24913 | 1,43305 | 701 |
| PDAC_Puleo_Endocrine | 0,046326 | 0,112726 | 0,26635 | -0,2164 | -1,1881 | 572 |
| PDAC_Puleo_Exocrine | 0,003018 | 0,011016 | 0,43171 | 0,25597 | 1,38764 | 355 |
| PDAC_Puleo_ICA9 | 3,61E-05 | 0,000264 | 0,55733 | 0,21065 | 1,27606 | #### |
| PDAC_Puleo_Immune | 0,013346 | 0,039765 | 0,38073 | -0,2177 | -1,2226 | 792 |
| PDAC_Puleo_StromaActiv | 3,35E-22 | 4,89E-20 | 1,22105 | -0,3825 | -2,1284 | 739 |
| PDAC_Puleo_StromaActivInflam | 0,000801 | 0,003545 | 0,47727 | -0,2382 | -1,34 | 845 |
| PDAC_Puleo_StromaInactive | 9,14E-08 | 1,21E-06 | 0,70498 | -0,274 | -1,5576 | 958 |
| PDAC_Puleo_Tech | 1,28E-06 | 1,33E-05 | 0,64355 | 0,23335 | 1,39285 | #### |
