## Supplementary Data 2 for "Favorable histo-molecular remodeling of pancreatic ductal adenocarcinoma after Total Neoadjuvant Therapy including Stereotactic Body Radiotherapy"

| pathway | pval | padj | log2err | ES | NES | size |
|---|---|---|---|---|---|---|
| CAF_FMG20_CAF.S1 | 0,7215 | 0,81035 | 0,0546 | -0,211 | -0,8637 | 72 |
| CAF_FMG20_IFNγ.iCAF | 0,3291 | 0,47494 | 0,1071 | 0,3677 | 1,1126 | 16 |
| CAF_FMG20_IL.iCAF | 0,0844 | 0,17365 | 0,2311 | 0,4093 | 1,3701 | 25 |
| CAF_FMG20_Normal.Fibroblast | 0,0002 | 0,00103 | 0,5188 | 0,4198 | 1,8386 | 80 |
| CAF_FMG20_TGFβ.myCAF | 0,1159 | 0,21984 | 0,1752 | -0,411 | -1,3368 | 25 |
| CAF_FMG20_detox.iCAF | 0,0001 | 0,00086 | 0,5188 | 0,6545 | 2,1018 | 21 |
| CAF_FMG20_ecm.myCAF | 0,0775 | 0,1688 | 0,2139 | -0,39 | -1,3929 | 36 |
| CAF_FMG20_wound.myCAF | 0,8946 | 0,91341 | 0,0456 | -0,209 | -0,717 | 31 |
| CAF_Neuzillet22_MYH11 | 0,9111 | 0,92377 | 0,0548 | 0,1953 | 0,6872 | 29 |
| CAF_Neuzillet22_PDPN | 0,4076 | 0,54096 | 0,0966 | 0,3151 | 1,0326 | 22 |
| CAF_Neuzillet22_POSTN | 0,3587 | 0,49668 | 0,1047 | 0,3059 | 1,0622 | 28 |
| CAF_Neuzillet22_subPOSTN | 0,3696 | 0,50427 | 0,101 | 0,3353 | 1,0586 | 19 |
| CAF_Turley20_hCAF0TGFB | 4E-08 | 7,2E-07 | 0,7195 | -0,463 | -2,1333 | 134 |
| CAF_Turley20_hCAF1early | 4E-06 | 4,3E-05 | 0,6105 | 0,3912 | 1,9283 | 148 |
| CAF_Turley20_hCAF2il6lif | 0,0666 | 0,15188 | 0,2878 | 0,234 | 1,252 | 258 |
| CAF_Tuveson19_iCAF | 4E-05 | 0,00031 | 0,5573 | 0,4443 | 1,9347 | 76 |
| CAF_Tuveson19_myo | 0,1025 | 0,20216 | 0,1919 | -0,49 | -1,3702 | 15 |
| CCCA_MetaProg_MP1.Cell.Cycle.G2.M | 0,0019 | 0,00749 | 0,4551 | -0,499 | -1,7829 | 36 |
| CCCA_MetaProg_MP10.Protein.maturation | 0,3189 | 0,47056 | 0,1133 | 0,2926 | 1,0967 | 38 |
| CCCA_MetaProg_MP11.Translation.initiation | 0,4829 | 0,60261 | 0,0881 | 0,2622 | 0,9827 | 38 |
| CCCA_MetaProg_MP12.EMT.I | 0,1783 | 0,30982 | 0,1373 | -0,324 | -1,2146 | 47 |
| CCCA_MetaProg_MP13.EMT.II | 0,0072 | 0,02405 | 0,407 | -0,431 | -1,6008 | 44 |
| CCCA_MetaProg_MP14.EMT.III | 0,3837 | 0,51877 | 0,101 | 0,2631 | 1,0401 | 45 |
| CCCA_MetaProg_MP15.EMT.IV | 0,5217 | 0,64554 | 0,0841 | 0,2473 | 0,9594 | 42 |
| CCCA_MetaProg_MP16.MES.glioma. | 0,0006 | 0,00269 | 0,4773 | -0,506 | -1,8484 | 42 |
| CCCA_MetaProg_MP17.Interferon.MHC.II.I. | 0,5273 | 0,64698 | 0,0699 | -0,262 | -0,9514 | 39 |
| CCCA_MetaProg_MP18.Interferon.MHC.II.II. | 0,1103 | 0,21198 | 0,2043 | 0,3344 | 1,278 | 41 |
| CCCA_MetaProg_MP19.Epithelial.Senescence | 0,0017 | 0,00687 | 0,4551 | 0,4686 | 1,7708 | 40 |
| CCCA_MetaProg_MP2.Cell.Cycle.G1.S | 0,3134 | 0,47056 | 0,0982 | -0,305 | -1,0897 | 36 |
| CCCA_MetaProg_MP20.MYC | 0,8851 | 0,91007 | 0,0568 | 0,1921 | 0,766 | 48 |
| CCCA_MetaProg_MP21.Respiration | 0,9839 | 0,98391 | 0,0528 | 0,1638 | 0,616 | 39 |
| CCCA_MetaProg_MP22.Secreted.I | 0,2815 | 0,44667 | 0,1221 | 0,2833 | 1,1195 | 46 |
| CCCA_MetaProg_MP23.Secreted.II | 0,0015 | 0,00643 | 0,4551 | 0,4493 | 1,7919 | 48 |
| CCCA_MetaProg_MP24.Cilia | 0,287 | 0,45055 | 0,1193 | 0,3246 | 1,1272 | 28 |
| CCCA_MetaProg_MP25.Astrocytes | 0,0433 | 0,10777 | 0,3218 | 0,3995 | 1,4331 | 33 |
| CCCA_MetaProg_MP26.NPC.Glioma | 0,7204 | 0,81035 | 0,0551 | -0,242 | -0,8418 | 32 |
| CCCA_MetaProg_MP27.Oligo.Progenitor | 0,1278 | 0,23565 | 0,1864 | 0,3751 | 1,3024 | 28 |
| CCCA_MetaProg_MP28.Oligo.normal | 0,8756 | 0,91007 | 0,0575 | 0,2112 | 0,7505 | 31 |
| CCCA_MetaProg_MP29.NPC.OPC | 0,8607 | 0,90401 | 0,0468 | -0,207 | -0,7543 | 41 |
| CCCA_MetaProg_MP3.Cell.Cylce.HMG.rich | 0,6443 | 0,75258 | 0,059 | -0,238 | -0,8853 | 43 |
| CCCA_MetaProg_MP30.PDAC.classical | 2E-05 | 0,00015 | 0,5756 | 0,5514 | 2,1644 | 44 |
| CCCA_MetaProg_MP31.Alveolar | 0,027 | 0,07792 | 0,3525 | 0,4311 | 1,5142 | 30 |
| CCCA_MetaProg_MP32.Skin.pigmentation | 0,3318 | 0,47494 | 0,111 | 0,3015 | 1,0774 | 32 |
| CCCA_MetaProg_MP33.RBCs | 0,0295 | 0,07969 | 0,3525 | 0,4139 | 1,4849 | 33 |
| CCCA_MetaProg_MP34.Platelet.activation | 0,315 | 0,47056 | 0,0982 | -0,316 | -1,0989 | 32 |
| CCCA_MetaProg_MP35.Hemato.related.I | 0,4022 | 0,53876 | 0,0972 | 0,293 | 1,0311 | 29 |
| CCCA_MetaProg_MP36.IG | 0,1732 | 0,30475 | 0,1563 | 0,3648 | 1,2565 | 27 |
| CCCA_MetaProg_MP37.Hemato.related.II | 0,7889 | 0,87041 | 0,0636 | 0,2204 | 0,8076 | 35 |
| CCCA_MetaProg_MP38.Glutathione | 0,2467 | 0,40929 | 0,1288 | 0,338 | 1,1457 | 26 |

| | | | | | | |
|---|---|---|---|---|---|---|
| CCCA_MetaProg_MP39.Metal.response | 0,1425 | 0,2569 | 0,1782 | 0,3234 | 1,236 | 41 |
| CCCA_MetaProg_MP4.Chromatin | 0,108 | 0,21017 | 0,1798 | -0,353 | -1,2925 | 42 |
| CCCA_MetaProg_MP40.PDAC.related | 0,5417 | 0,65903 | 0,0827 | 0,2424 | 0,9514 | 44 |
| CCCA_MetaProg_MP41.Unassigned | 0,0841 | 0,17365 | 0,2043 | -0,385 | -1,367 | 35 |
| CCCA_MetaProg_MP5.Stress | 0,0272 | 0,07792 | 0,3525 | 0,374 | 1,4779 | 46 |
| CCCA_MetaProg_MP6.Hypoxia | 0,4333 | 0,56488 | 0,0798 | -0,269 | -0,9984 | 44 |
| CCCA_MetaProg_MP7.Stress.in.vitro. | 0,08 | 0,17176 | 0,2378 | 0,393 | 1,3829 | 29 |
| CCCA_MetaProg_MP8.Proteasomal.degradatio | 0,229 | 0,38881 | 0,1365 | 0,2916 | 1,1561 | 47 |
| CCCA_MetaProg_MP9.Unfolded.protein.respo | 0,6905 | 0,79378 | 0,0715 | 0,2255 | 0,8744 | 43 |
| CCK_STIM_Desert-like | 0,8549 | 0,90401 | 0,0587 | 0,1971 | 0,7817 | 47 |
| CCK_STIM_Hepati_Stem-like | 0,25 | 0,41011 | 0,1143 | -0,287 | -1,133 | 60 |
| CCK_STIM_Immune_Classical | 0,4526 | 0,57328 | 0,0775 | -0,267 | -0,9901 | 44 |
| CCK_STIM_Inflammatory_Stroma | 0,0935 | 0,18965 | 0,1958 | -0,329 | -1,3152 | 66 |
| CCK_STIM_Tumor_Classical | 0,5583 | 0,66275 | 0,068 | -0,235 | -0,9367 | 64 |
| CCK_Sia13_INFLAMGenes | 0,2726 | 0,44216 | 0,1084 | -0,271 | -1,1161 | 78 |
| CCK_Sia13_PROLIFGenes | 0,0086 | 0,02805 | 0,3807 | -0,214 | -1,2151 | 1080 |
| DrugBank_Fluorouracil | 0,3128 | 0,47056 | 0,1124 | 0,4089 | 1,1249 | 12 |
| DrugBank_Gemcitabine | 0,8475 | 0,90401 | 0,0559 | 0,2748 | 0,7163 | 10 |
| DrugBank_Irinotecan | 0,8496 | 0,90401 | 0,0558 | 0,2739 | 0,7138 | 10 |
| DrugBank_Oxaliplatin | 0,0011 | 0,00512 | 0,4551 | 0,7661 | 1,9432 | 9 |
| DrugBank_Paclitaxel | 0,7929 | 0,87041 | 0,0531 | -0,281 | -0,7338 | 11 |
| ECM_Helms22_PSCcaf | 4E-06 | 4,3E-05 | 0,6105 | -0,474 | -2,0194 | 90 |
| IMMU_GenJCI121924_GeneralImmu | 0,3191 | 0,47056 | 0,1396 | 0,1845 | 1,038 | 482 |
| IMMU_GenJCI121924_ICKrelated | 0,6147 | 0,72373 | 0,0644 | -0,319 | -0,8909 | 14 |
| IMMU_MCPcounter_B.lineage | 0,3157 | 0,47056 | 0,1093 | 0,4877 | 1,1476 | 7 |
| IMMU_MCPcounter_CD8Tcells | 0,2174 | 0,3734 | 0,1336 | 0,8904 | 1,1955 | 1 |
| IMMU_MCPcounter_Cytotox.lymph | 0,4524 | 0,57328 | 0,0829 | -0,599 | -1,0346 | 3 |
| IMMU_MCPcounter_Endothelial | 0,0605 | 0,14014 | 0,3218 | 0,4593 | 1,4705 | 20 |
| IMMU_MCPcounter_Fibroblasts | 0,0704 | 0,15806 | 0,2311 | -0,623 | -1,4603 | 8 |
| IMMU_MCPcounter_Mono.lineage | 0,5547 | 0,66275 | 0,0708 | -0,417 | -0,9415 | 7 |
| IMMU_MCPcounter_Myeloid.dendritic | 0,4286 | 0,56371 | 0,0894 | 0,7713 | 1,0357 | 1 |
| IMMU_MCPcounter_NK | 0,5487 | 0,66201 | 0,0761 | 0,7214 | 0,9687 | 1 |
| IMMU_MCPcounter_Neutrophils | 0,8278 | 0,89523 | 0,0508 | -0,308 | -0,7227 | 8 |
| IMMU_MCPcounter_Tcells | 0,7115 | 0,81035 | 0,0596 | -0,387 | -0,789 | 5 |
| PDAC_Bailey16_ADEX | 0,0033 | 0,01139 | 0,4317 | -0,368 | -1,6079 | 98 |
| PDAC_Bailey16_Immunogenic | 0,2757 | 0,44233 | 0,125 | 0,2561 | 1,1047 | 74 |
| PDAC_Bailey16_Progenitor | 0,1178 | 0,22046 | 0,1938 | 0,3745 | 1,3179 | 29 |
| PDAC_Bailey16_Squamous | 6E-05 | 0,00045 | 0,5573 | -0,291 | -1,5584 | 449 |
| PDAC_CSY20_BasallikeA | 0,049 | 0,11914 | 0,3218 | 0,2343 | 1,2473 | 245 |
| PDAC_CSY20_BasallikeB | 0,0253 | 0,07532 | 0,3525 | -0,324 | -1,4158 | 99 |
| PDAC_CSY20_ClassicalA | 8E-06 | 7,7E-05 | 0,5933 | 0,3438 | 1,8134 | 218 |
| PDAC_CSY20_ClassicalB | 0,0335 | 0,08742 | 0,3218 | 0,2962 | 1,3445 | 96 |
| PDAC_CSY20_Sig11 | 0,4555 | 0,57328 | 0,0759 | -0,225 | -1,0031 | 112 |
| PDAC_CSY20_Sig12 | 0,0592 | 0,13949 | 0,245 | -0,291 | -1,3021 | 118 |
| PDAC_CSY20_Sig3 | 0,0373 | 0,09555 | 0,282 | -0,226 | -1,2191 | 497 |
| PDAC_CSY20_Sig4 | 0,3591 | 0,49668 | 0,1115 | 0,2114 | 1,0363 | 140 |
| PDAC_CSY20_Sig5 | 0,0305 | 0,08087 | 0,3525 | 0,2174 | 1,233 | 515 |
| PDAC_CSY20_Sig7 | 0,3485 | 0,49403 | 0,1183 | 0,1984 | 1,0408 | 213 |
| PDAC_CSY20_Sig8 | 0,0282 | 0,07903 | 0,3525 | 0,2385 | 1,278 | 255 |
| PDAC_CSY20_Sig9 | 0,1023 | 0,20216 | 0,2139 | 0,3483 | 1,3161 | 40 |

| | | | | | | |
|---|---|---|---|---|---|---|
| PDAC_Hwang22_Fibro.Adhesive | 0,0002 | 0,00096 | 0,5188 | -0,351 | -1,6956 | 180 |
| PDAC_Hwang22_Fibro.Immunomodulatory | 0,0026 | 0,00967 | 0,4317 | 0,3065 | 1,517 | 153 |
| PDAC_Hwang22_Fibro.Myofibroblastic | 2E-13 | 7,1E-12 | 0,9326 | -0,498 | -2,3985 | 176 |
| PDAC_Hwang22_Fibro.Neurotropic | 0,0114 | 0,03623 | 0,3807 | 0,2724 | 1,3711 | 169 |
| PDAC_Hwang22_Malignlineage.Acinarlike | 2E-05 | 0,00015 | 0,5756 | 0,36 | 1,7984 | 162 |
| PDAC_Hwang22_Malignlineage.Basaloid | 3E-06 | 3,9E-05 | 0,6273 | 0,3729 | 1,873 | 166 |
| PDAC_Hwang22_Malignlineage.Classicallike | 0,1491 | 0,26547 | 0,1847 | 0,2261 | 1,1429 | 173 |
| PDAC_Hwang22_Malignlineage.Mesenchymal | 0,6852 | 0,79378 | 0,0525 | -0,189 | -0,9143 | 178 |
| PDAC_Hwang22_Malignlineage.Neurallikeprog | 0,3606 | 0,49668 | 0,0871 | -0,223 | -1,0444 | 153 |
| PDAC_Hwang22_Malignlineage.Neuroendocri | 0,8196 | 0,89298 | 0,0478 | -0,19 | -0,8322 | 98 |
| PDAC_Hwang22_Malignlineage.Squamoid | 0,0436 | 0,10777 | 0,3218 | 0,2571 | 1,2796 | 158 |
| PDAC_Hwang22_Malignstate.Adhesive | 0,966 | 0,97263 | 0,0376 | -0,16 | -0,7619 | 167 |
| PDAC_Hwang22_Malignstate.CyclingG2M | 3E-07 | 4E-06 | 0,675 | -0,44 | -2,0331 | 136 |
| PDAC_Hwang22_Malignstate.CyclingS | 0,018 | 0,05489 | 0,3525 | -0,312 | -1,4476 | 141 |
| PDAC_Hwang22_Malignstate.Interferonsignali | 0,0002 | 0,00098 | 0,5188 | -0,364 | -1,707 | 152 |
| PDAC_Hwang22_Malignstate.MYCsignaling | 0,084 | 0,17365 | 0,1958 | -0,256 | -1,2281 | 172 |
| PDAC_Hwang22_Malignstate.Ribosomal | 5E-20 | 7,8E-18 | 1,1602 | 0,544 | 2,782 | 182 |
| PDAC_Hwang22_Malignstate.TNFNFkBsignalin | 9E-09 | 1,7E-07 | 0,7477 | -0,435 | -2,0796 | 169 |
| PDAC_Moffitt15_ActivatedStroma | 2E-13 | 7,1E-12 | 0,9326 | -0,348 | -1,9156 | 646 |
| PDAC_Moffitt15_BasalLike | 3E-10 | 5,4E-09 | 0,814 | -0,348 | -1,8667 | 469 |
| PDAC_Moffitt15_CellCycle | 0,0001 | 0,00085 | 0,5188 | -0,22 | -1,2917 | 2170 |
| PDAC_Moffitt15_Classical | 0,0003 | 0,00176 | 0,4985 | 0,2396 | 1,385 | 618 |
| PDAC_Moffitt15_Endocrine | 0,4383 | 0,56626 | 0,1124 | 0,1818 | 1,0047 | 397 |
| PDAC_Moffitt15_Exocrine | 0,0002 | 0,00103 | 0,5188 | -0,289 | -1,5265 | 393 |
| PDAC_Moffitt15_F10 | 0,0564 | 0,13506 | 0,2165 | -0,204 | -1,1563 | 1088 |
| PDAC_Moffitt15_F14 | 0,0021 | 0,00793 | 0,4317 | 0,2021 | 1,2211 | 1108 |
| PDAC_Moffitt15_F2 | 2E-10 | 5,4E-09 | 0,8267 | 0,2702 | 1,6162 | 981 |
| PDAC_Moffitt15_Immune | 0,1291 | 0,23565 | 0,1455 | -0,204 | -1,1161 | 627 |
| PDAC_Moffitt15_Liver | 0,2446 | 0,40929 | 0,1563 | 0,1914 | 1,0598 | 409 |
| PDAC_Moffitt15_Lung | 0,0004 | 0,00208 | 0,4985 | 0,2951 | 1,5582 | 233 |
| PDAC_Moffitt15_Muscle | 0,0129 | 0,03993 | 0,3807 | 0,2162 | 1,2391 | 582 |
| PDAC_Moffitt15_NormalStroma | 0,0007 | 0,00315 | 0,4773 | 0,2295 | 1,3329 | 673 |
| PDAC_PDAssigner_Classical | 0,0017 | 0,00687 | 0,4551 | 0,5862 | 1,9207 | 22 |
| PDAC_PDAssigner_Exocrine | 0,0028 | 0,01017 | 0,4317 | -0,591 | -1,8002 | 20 |
| PDAC_PDAssigner_QM | 0,0066 | 0,02231 | 0,407 | -0,629 | -1,7584 | 15 |
| PDAC_PDXph1_basal | 0,003 | 0,01085 | 0,4317 | -0,239 | -1,3181 | 663 |
| PDAC_PDXph1_classical | 0,0003 | 0,00176 | 0,4985 | 0,254 | 1,4332 | 477 |
| PDAC_Puleo_Basal | 3E-14 | 1,2E-12 | 0,976 | -0,357 | -1,957 | 618 |
| PDAC_Puleo_Classic | 4E-07 | 5,6E-06 | 0,675 | 0,2638 | 1,5419 | 701 |
| PDAC_Puleo_Endocrine | 0,0287 | 0,07903 | 0,3525 | -0,227 | -1,2367 | 572 |
| PDAC_Puleo_Exocrine | 0,0015 | 0,00643 | 0,4551 | -0,269 | -1,4029 | 355 |
| PDAC_Puleo_ICA9 | 0,8834 | 0,91007 | 0,0285 | -0,158 | -0,916 | 1736 |
| PDAC_Puleo_Immune | 0,0769 | 0,1688 | 0,1864 | -0,206 | -1,1487 | 792 |
| PDAC_Puleo_StromaActiv | 7E-19 | 5,4E-17 | 1,1239 | -0,373 | -2,0678 | 739 |
| PDAC_Puleo_StromaActivInflam | 0,3258 | 0,47494 | 0,1483 | 0,1729 | 1,0273 | 845 |
| PDAC_Puleo_StromaInactive | 0,7311 | 0,81484 | 0,0412 | -0,167 | -0,9418 | 958 |
| PDAC_Puleo_Tech | 0,0001 | 0,00074 | 0,5384 | 0,2128 | 1,2992 | 1330 |
